# ‘Sniff Olfactometer (SO) Protocols

**DOI:** 10.1101/2022.07.08.499357

**Authors:** Jiayue Ni, Qi Tang, Jianbo Dave Huang, Leto Solla, Hannah Kelson, Marcus Weeks, Zoe Alcott, Justin Ong, Andrea Gomez, Kaifeng Ding, A. Terry E. Acree

**Affiliations:** Cornell University

## Abstract

Most olfactometers used to study human olfaction have stimulus durations of more than 1 second and often lasting minutes(Dravnieks 1975; Leland *et al*. 2001; Schmidt and Cain 2010). During long stimulations, olfactory receptor responses and their resulting behaviors are modulated by adaptation and habituation to the stimulus(Pellegrino *et al*. 2017; Rankin 2009; Wilson and Linster 2008). For example, EOG results from the first deorphanized olfactory receptor tissue reached a maximum in ∼1 s, dropping to 1/2 maximum in the next second, and showing little signal reduction until the stimulation stopped after 6 seconds(Zhao *et al*. 1998). Longer stimulations can result in complete habituation; receptors still respond even though the behavior shows complete habituation (Barwich 2014). To minimize the effects of adaption and habituation on stimulus responses, the sniff olfactometer (SO) combined the precision of a blast olfactometer with the gentleness of a stream olfactometer by blasting a brief odorant puff (70ms duration) into a subject’s self-imposed inhalation air stream(Rochelle 2017; Rochelle *et al*. 2017b; Wyckoff and Acree 2017). Here we describe SO protocols for threshold determinations of odorants in aqueous headspaces using odorant recognition probabilities associated with Log(odorant-concentrations(Rochelle *et al*. 2017a)). During a single trial a subject, preconditioned to associate a veridical name with a given odor (e.g., a pyrazine with “nuts” when the odor was detected and “not nuts” if it wasn’t) was cued to “inhale” and 750ms later, a 15ml-70ms puff of odorant headspace was delivered into their inhalation airstream. A session consisted of 12 randomized double-blind trials of 3 different odorant concentrations. Additional sessions with different concentrations were conducted until the response probability to the samples ranged from below 0.2 to above 0.8. The robustness of the fitted function and the size of their confidence intervals depended on the difference between the concentrations of the odorants during a single session: small differences in sample concentration resulted in the data failing to fit a logistic function; larger concentration differences resulted in a better fit to the model. However, if one of the stimuli had no odorant at all *i*.*e*., a blank, the response to the blank was random.

## Introduction

It is generally acknowledged that humans can recognize odor mixtures as either a single percept (a configuration) or as individual component-odors (elements) experienced simultaneously (Pellegrino *et al*. 2017; Sinding *et al*. 2013; Sinding *et al*. 2011; Thomas-Danguin *et al*. 2014). How an animal experiences and responds is driven by an attentional top-down pattern recognition strategy and the salience of the stimulation event to the organism(Romagny *et al*. 2018). This process is shared with many other animals, despite millions of years of evolutionary separation. Human odor recognition, caused by odorants that smell fruity, meaty, bloody, smokey, etc., provide humans with clues to changes in their environment, the quality of their food, or warn them of danger or the possibility of pleasure. An essential tool used for the last 150 years to study the process of olfaction are olfactometers of many different designs.

Many olfactometers were designed to deliver stimulants to e.g. insects and an assortment of other animals, measuring changes in behavior, physiology, or neurobiology(Laing *et al*. 1994; Wilson and Stevenson 2006). These experiments advanced our understanding of olfaction in non-human models, but it told us very little about human perception and the emotional responses that govern human olfactory choices. Previous studies with humans generally related odorant compositions to semantic labels based on detection, recognition, or scaling(Lawless and Heymann 2010).

Unfortunately, scaling—a cross-modal matching procedure—requires more computational power than either detection or recognition, resulting in very noisy data. However, if we make measurements in humans that parallel the olfactometry and cognitive measures used with non-human subjects, we can develop insights from human research that are applicable to understanding animal behavior and *vice-versa(Wilson and Stevenson 2006)*. Among the drawbacks of existing human olfactometers and protocols are: 1) irreproducible stimulant compositions (unless equilibrium headspace s are sampled), 2) adaption and habituation due to long stimulant exposure times, 3) plumbing contamination during the experiment and 4) scaling-induced noise. This paper describes a Sniff Olfactometer (SO)(Wyckoff and Acree 2017) and protocols that optimize inter-stimulus intervals, and noticeable differences for the study of unary and binary mixture perceptions.

## Protocol

Because smell-evoked behavior in humans is dependent on prior experiences, memories of events, and their emotional tags, appropriate human conditioning is essential for meaningful experimental designs(Cain 1979). To optimize conditioning, minimize habituation, and quantify stochasticity, the SO synchronizes each inhalation with double-blind stimulation and forced-choice paradigms. The threshold measurement over 6 concentrations for one subject took less than 15 minutes while each trial and session took less than 10 seconds and 6 minutes respectively on the SO^9-11^. Conditioning trials were prior to the threshold measurement session to familiarize the odorant in the experimental context. This enhanced a subject’s ability to associate the odorant with the language context, in this case, the veridical name assigned to an odorant. Subjects who have been well-conditioned perform with greater precision and accuracy on identification tasks, compared to unconditioned or naÏve subjects (Cain and Algom 1997). In each session, 12 trial responses were expressed as probabilities (≤ 1.0) of perceiving the odorant, e.g., P(“nut”) = (number of “nut” responses) / (total number of puffs) were fitted to a logistic binomial regression, i.e., P(“nut”) = f(Log[odorant])(Rochelle *et al*. 2017a).

The SO used in this experiment (Figure 1), was derived from the instrument patented in 2018(Rochelle *et al*. 2017b; Wyckoff and Acree 2017). Aqueous samples (50mls 10% PEG) were placed in 250ml Fluro-ethylene polymers (FEP) bottles and equilibrated on a shaker table for at least 1hr but less than 24hrs before the experiment. Per session, each bottle containing three different odorant concentrations was mounted to the sniff block; this created a stimulus triad, with each stimulus traveling via separate tubes (<3 cm) to the sniff port (Figure 1B). Subjects sat in front of the SO, positioned their heads with the aiming system, and placed one hand on the mouse to operate the computer (Figure 1A). Instructing cues on the monitor guide d subjects to aim and inhale (Figure 1B). 700ms after the inhale cue, the actuator delivered 15mL of the odorant headspace mixture into the inhalation stream. Subjects were then cued to answer a binary question displayed on the monitor regarding the veridical name of the stimulus presented. A binary choice was required to proceed to the next trial.

**Figure 1.**
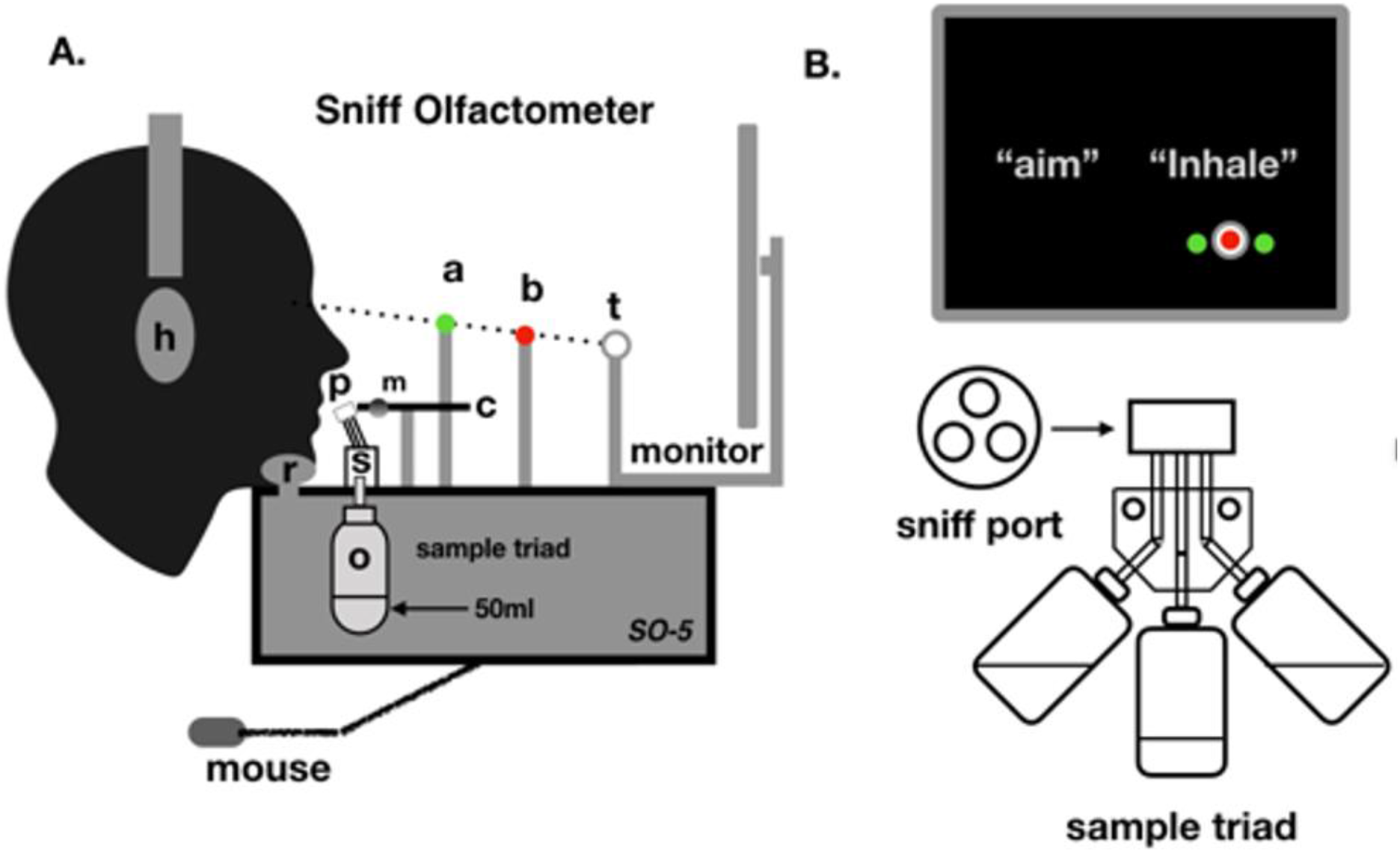
(A) Side view of the SO-5. headphones **h**, chin rest **r**, sniff port **p**, audio detector **m**, and **c** aiming system using **a, b**, and **t** to fix the head position, (B) Instructing cues showing on the monitor and the stimulant triad used in a single session. Ding et al., (2021)(Ding *et al*. 2019)

The SO-5 was controlled by PsychoPy^©^, a Python-based computer program to provide precise timing for all visual and auditory cues, data collection, and data storage(Peirce 2022; Peirce *et al*. 2019; Peirce 2007) (Figure 1). To eliminate any sound disturbances from the surroundings, subjects listen to their own music in their headphones.

### Preliminary results from SO measurements

In 2017, Rochelle et al.(Rochelle 2017; Rochelle *et al*. 2017a) developed and refined the threshold protocol used to determine a subjects’ equal odds ratio (EOR), the ratio of concentrations of components in a two-odorant mixture that led to identical probability of perceiving either of the components. In preliminary work, they observed that presenting the stimulants in increasing or decreasing concentration—as would be the case in a classical staircase method(Cornsweet 1962)—the data was extremely noisy, yielding estimates with large confidence intervals. However, by alternating or interlacing the concentrations, more robust data was obtained. In early 2020, Qi Tang and Jiayue Ni attempted to investigate this interlacing effect on subject performance (Figure 2). Using the SO-5, thresholds for hexanal with 12 concentrations were performed, using both the traditional staircase model (‘sequential’), and the interlaced method noticed by Rochelle et al (‘alternate’). Due to COVID-19, a full trial was not permissible; however, preliminary data collected prior to shutdown showed the increased subject performance using the ‘alternate’ method noted by Rochelle et al, when compared to the conventional ‘sequential’ method (Figure 2).

**Figure 2.**
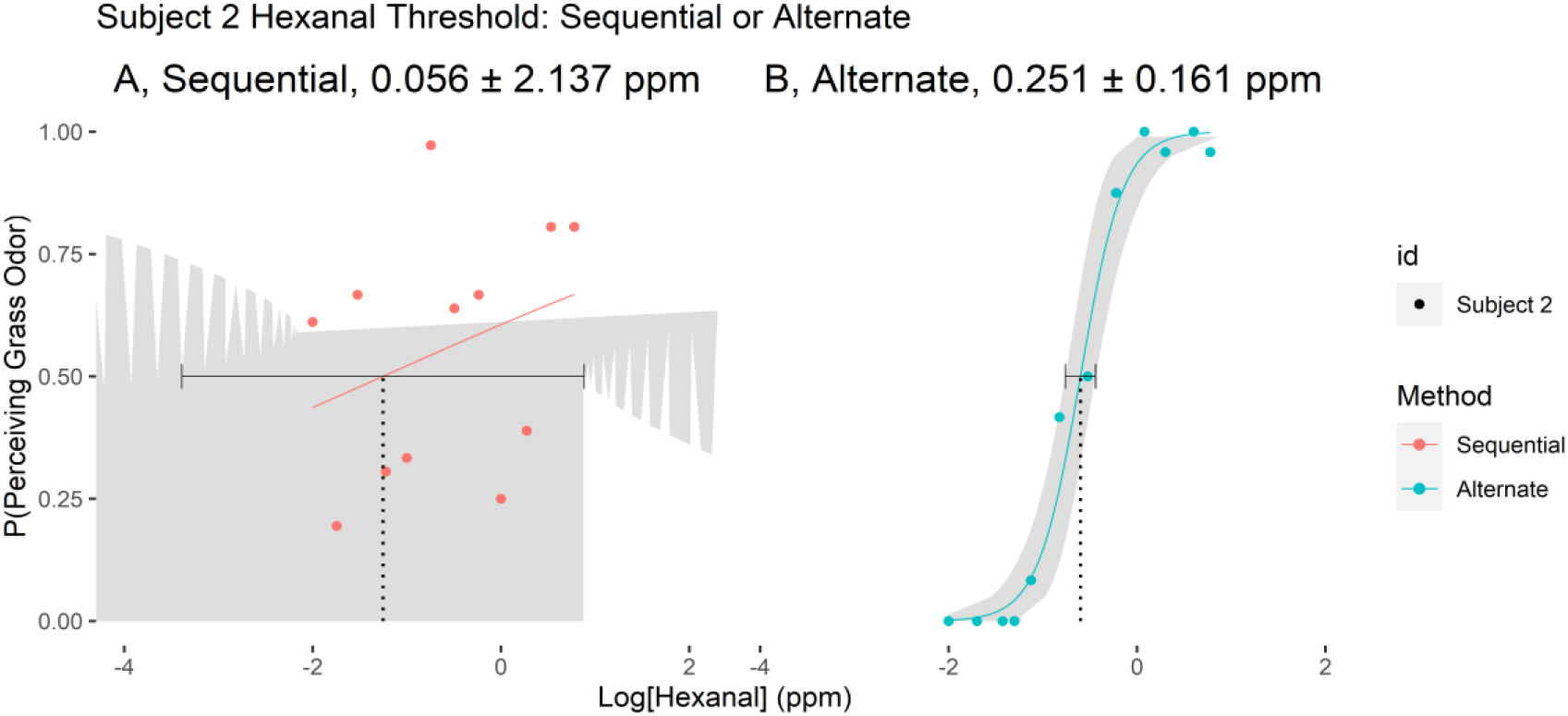
Plots of hexanal recognition probability P(“grass”) using a sequential vs alternated presentation for one subject. The x-axis is the log10 of the sample concentrations (in parts/million), and the y-axis is the probability the subject chose “grass” or “blank” after stimulation. The error bars are the CIs at 0.95% (Tang 2020 unpublished). This study was terminated in April 2020 due to COVID-19.

### New SO Protocol

In January 2021, when research was again allowed at the University, we compared the sequential (SEQ) vs alternated (ALT) presentation using 7 subjects and two odorants: hexanal and 2,3,5-trimethylpyrazine.

In the preliminary work done by Tang and Ni before the COVID-19 shutdown, each concentration (n = 12) was puffed 36 times, leading to the subject being required to sniff and identify the compound 432 times. When the group replicated the experiment, it was determined that the whole process took 45 - 80 minutes/subject, and according to the subjects, “they lost focus and felt fatigue after about 30 minutes”.

We therefore modified the protocols to require 6 concentrations, with each concentration being puffed 12 times, meaning that each threshold measurement would be the result of 72 total puffs. This change in protocol reduced the time needed to collect each threshold to around 30min. Air in the test room was exchanged 6 times/hr. and filtered to remove undesired odorants(Delahunty *et al*. 2006; Foster *et al*. 1950); additionally, all subjects and researchers were asked not to use any highly fragranced shampoos or soaps and to avoid smoking, drinking, eating and similar activities for at least 1 hour before the experiment(Goyert *et al*. 2007; Wise *et al*. 2000).

### Subjects

2 Male and 5 female subjects were chosen from the students allowed to be in the building; this restriction was due to university regulations regarding COVID-19. Some subjects had experience with the SO previously, but none of the subjects knew any of the experimental details before the project was completed. They were asked to participate in two sessions per experiment and to give the researchers feedback as to how they felt about the experimental session (if they were tired, etc.), and other comments they wanted to share about the experiment, the equipment, and the researcher. They were also encouraged to ask questions but were told that some questions would not be answered by the researcher. The survey results indicated that all the subjects used in the experiment reported they could detect the target odors during the entire experiment, and that it was not fatiguing. All subjects were conditioned first by a “teaching session” and then by a brief priming with the stimulus immediately before the “testing session”.

### Chemical Stimuli

Because of the poor solubility of most odorants in water, pure stimulants were first dissolved in polyethylene glycol (PEG). Aliquots of these PEG stock solutions were dispersed into water and diluted until a target concentration was achieved. The stimuli produced by the SO-5 were 15 ml headspace puffs of **hexanal** (HEX, the veridical name “green”) (CAS Registry No. 66-25-1, >98%) or **2**,**3**,**5-trimethylpyrazine** (TMP, the veridical “nut”) (CAS Registry No. 14667-55-1, >99%) diluted in 10% polyethylene glycol (PEG, veridical name “blank”) (CAS Registry No. 9002 -88-4) prepared in deionized water from the 400 ppm (parts/million) stock solution. HEX and TMP were selected for their dissimilar chemistry and odor(Wise *et al*. 2000). All solutions were prepared from stock solutions within 4 to 12 hours before the first session and stored in amber bottles at room temperature (22 °C).

### Stimulus Presentation

Thresholds were determined for each subject from 6 different concentrations ranging from (HEX: 0.01ppm to 15ppm and TMP: 0.1ppm to 20ppm) depending on their individual sensitivity to the stimulus. A veridical name was assigned to each stimulant by the experimenter; “green” for HEX, “nut” for TMP. The subject put their chin on the chin rest to control their head from movement; correct position was ensured by directing the subject to sight down an aiming process, which positioned the head in the correct position over the sniff block. The monitor would instruct the subjects to follow the experimental process, e.g., “click when you are ready”, “inhale”, and “select what you just smelled”. Subjects were then presented with a binary forced choice test, where the options were either the veridical name of the compound, or “blank”, if the subject smelled nothing in the puff. All puffs presented to the subject contained the compound being tested in the threshold. This presentation process was repeated until each bottle in the trio was presented to the subject 4 times, at which point the bottles were rearranged to avoid association between a particular concentration and any positional difference in the SO. Each bottle was present in each position of the sniff block during a given test of a trio of concentrations, allowing for 12 presentations of each compound; this allowed for a 36 -puff session, in which the order of presentation was double-blind. After 36 total puffs, the subject was then given a 2-5-minute break, in which the experimenter exchanged the first trio and sniff block for a second trio and sniff block containing the remaining 3 concentrations for the test.

### Subject Conditioning

Each new subject was given a detailed explanation of the human-subjects consent form (Appendix) to ensure they understand what their rights and responsibilities were. Then high concentrations, 20-50 ppm of the target odorants, were presented to condition the subjects to become familiar with and learn to distinguish them. Each teaching session used a triad of odorant solutions (50ms each) sometimes including a 10% PEG (blank) installed in the SO sniff block. A pre-programed PsychoPy^©^ script displayed instructions on the monitor e.g., “click to start”, “get ready”, “Inhale”, “what did you smell?”. Also displayed on the screen with “What did you smell?” was “nut”, “green”, or “blank” depending on to sample puffed; the veridical name displayed always corresponded correctly to the presented stimulus, eg. “nut” for TMP, “green” for HEX, and “blank” for 10% PEG. The only difference between a teaching session and a testing session was the inclusion of the name of the target compound along with the question “What did you smell?”.

Each bottle would be puffed 5 times and each time the subjects were asked to identify the smell by associating it with one of the veridical names indicated on the screen. The subjects were trained to distinguish among nutty, green, and the blank smell as well as to become familiar with the SO system and to experience a typical session. If a subject reported that it was hard for them to perceive the smell of the puffs, the concentration range would shift up one and the entire teaching session would be done again. Subjects that attained 13 out of 15 correct (13/15) were allowed to move to the next step, whereas the ones that failed this screening were required to take the teaching again until they reached a 13/15 correctness. The subjects who failed to achieve 13/15 correctness within 5 teaching sessions of the teaching-testing cycle were not used with the project. All subjects were required to take the test again before each formal experiment, i.e., if the experiments were done over more than one day, the subjects were required to take the test the next day.

### Testing Session

During testing, fresh preparations of the samples used in training (HEX and TMP, in 10% PEG) were prepared, spread exponentially over a range of concentrations (HEX: 0.01ppm to 15ppm; TMP: 0.1ppm to 20ppm). A single trial, ∼7s in duration, began with a mouse click by the subject followed in 1 second by the instruction to inhale deeply and at a moderate speed; 750 ms later the SO puffed a 15ml sample (70ms duration) of the headspace from one of the bottles into the inhalation-air stream. After ∼1500ms of the first cue the subject was asked to make a binary choice, between “blank” and either “green” or “nut” depending on the chemical being tested. Following this choice, the next trial would begin. After each bottle in the triad was puffed 4 times, the bottle positions were switched resulting in 12 repeats for each concentration of an odorant solution rotated through each bottle position. A session (36 trials) consisted of the 4 trials at each bottle of the 3 bottle-positions for each of the 3 concentrations tested. Between each 36-trial session, subjects were given a rest period of around 5 minutes.

## Experiments

### Threshold measurements

Because odor thresholds are highly idiosyncratic ranging over orders of magnitude, we began the study by determining the thresholds for each subject and each compound. We then prepared stimuli for each subject that included their threshold near the middle of the range. Depending on the presentation order being tested (SEQ or ALT), a different bottle order was used in each triad (Figure 4). Only the bottle arrangement for the first session is shown, the second session was similar but included bottles #3, #4, and #5 for the sequential (SEQ) condition, and bottles #2, #4, and #6 for the alternate (ALT) condition. Once a subject successfully completed training, they were presented a triad of bottles as described above and a PsychoPy script executed followed by bottle-rotation before the next session. After three sessions, new bottles at 3 new concentrations were used according to the order shown (Figure 4).

**Figure 3.**
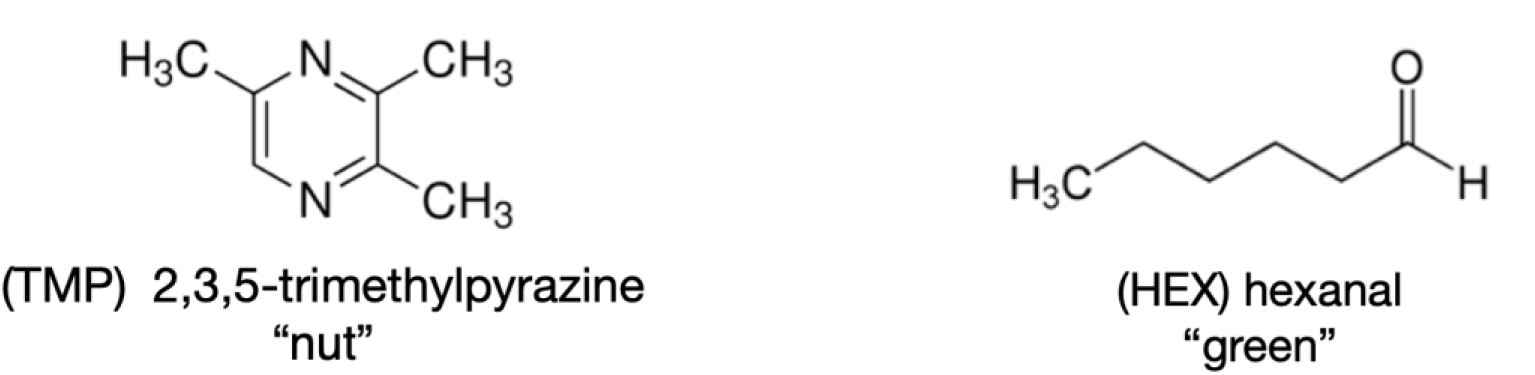
Chemical Structures of hexanal (HEX) and 2,3,5-trimethylpyrazine (TMP). After discarding the solutions, all bottles and caps were cleaned with 6 times with deionized water and once with 70% ethanol. If the bottles and caps still retained odor after this washing process, they were discarded.

**Figure 4.**
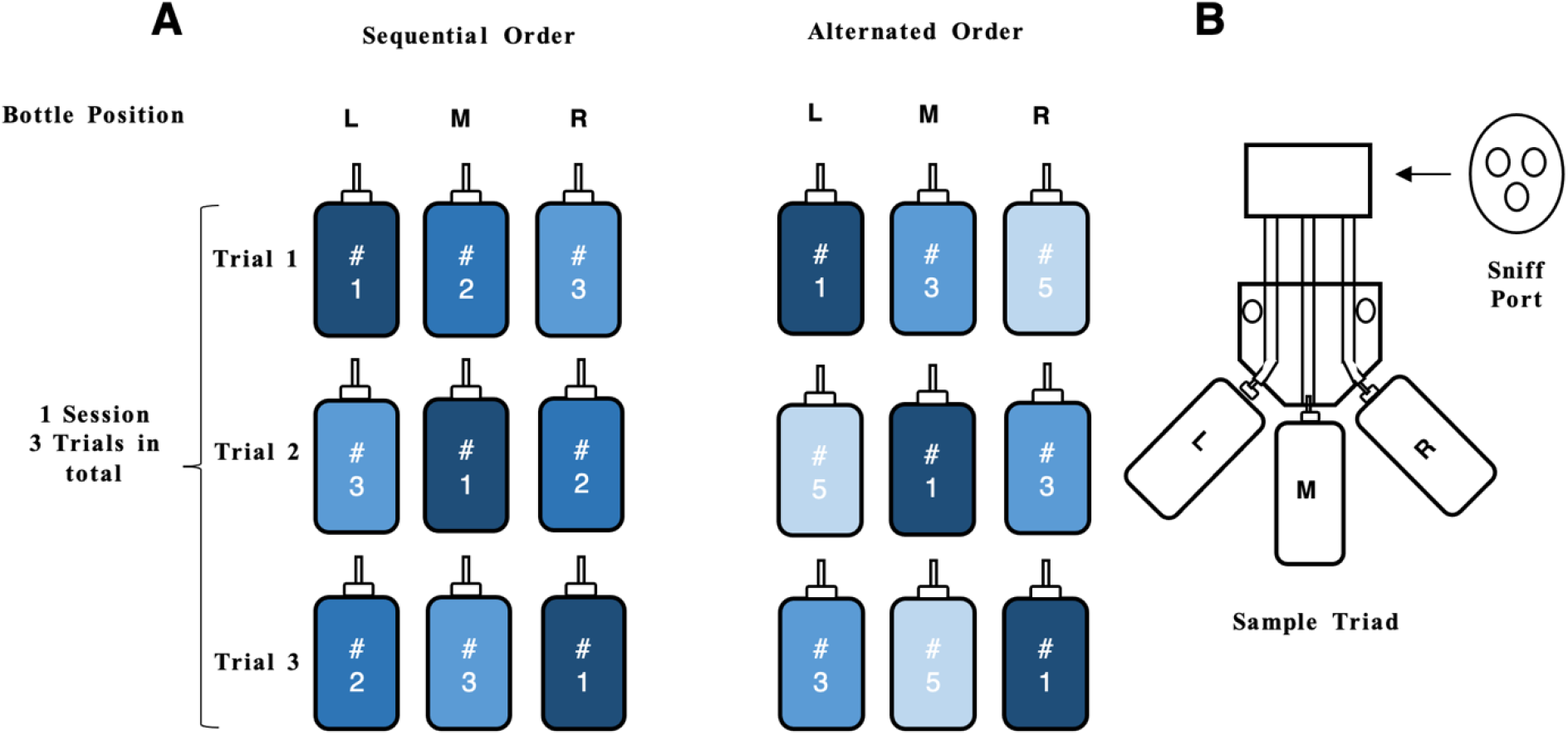
A. Example of sequential and alternated orders of bottles #1, #2, #3 within a session and the changes when the bottles were switched after each trial and B. Bottles in the sample triad. L (left position), M (middle position), and R (right position). The darker the color, the more concentrated the sample solution.

### Reproducibility

Six subjects (B-G) repeated the experiments 2-3 days in a row with fresh solutions. Subject A repeated the threshold measurements on 3 separate days using the same solution used on the first day. Only subject A participated in this test with recycled samples, while the other subjects (B -G) took these tests with fresh sample solutions.

### With a blank vs without a blank

Subjects A, B, and C had thresholds that included a blank of pure PEG without an odorant added as one of the 6 offered bottles. Subjects D, E, F, and G had thresholds without a blank (ie; all 6 bottles presented in a threshold had an odorant present). This experiment was done to see what the effect of a blank would be on the threshold, as well as on model fit.

### Statistical Analysis

All data were analyzed with R(Team 2020). All word responses, i.e., “nutty” and “blank” (Figure 9), were converted to binary values, i .e., 1 and 0. Averages were taken based on these numbers by concentration. Log 10 of concentrations were used to make the Gaussian curves (all curves are in the Appendix). In each data set, there are 6 points (one for each sample), each of which was the average of 12 observations by each subject, i.e., the completion of one group of experiments. The points generalized the linear models which were analyzed by MASS package(Herz *et al*. 2022; MASS 2020) in R to create dose response curves. As each subject had multiple measurements of their threshold per chemical, boxplots were created to visualize the variance between a subject’s threshold across the days of testing (all boxplots can be found in the Appendix). These boxplots were based upon the threshold points and the confidence intervals extracted from the curves.

## RESULTS

### Sequential VS Alternated Order

A threshold curve for one subject responding to the SEQ and ALT presentations of TMP is shown in Figure 5. The fit of the data to a binomial logistic function is more precise for alternate presentation when compared to sequential presentation; the obtained thresholds are not significantly different from each other, while the confidence interval produced by the alternate presentation is markedly narrower than the sequential confidence interval.

**Figure 5.**
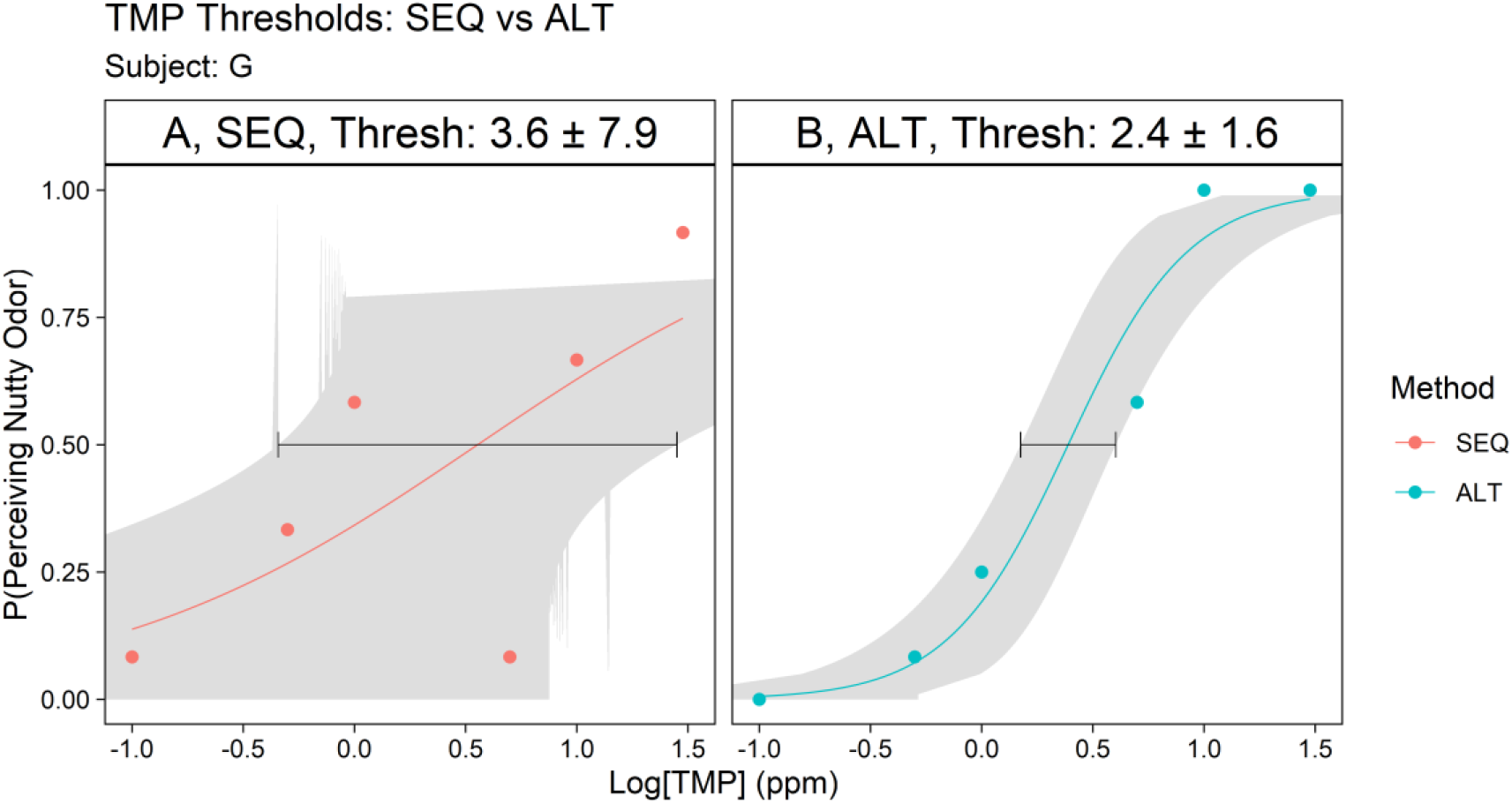
TMP threshold measurement, Subject G, replication 1. The samples were freshly made. The x-axis is the log10 of the sample concentrations (in ppm), and the y-axis is the probability the subject perceived a sample and identified it as “nut”. For TMP, 20, 10, 5, 1, 0.5, and 0.1 ppm were chosen as concentrations.

Figure 6 displays the curves generated for a subject experiencin g the SEQ and ALT presentations of HEX, with the addition of a 10% PEG blank. In the sequential presentation, the addition of a 10% PEG blank substantially disrupted the fit and confidence of the model, an effect that was mitigated in the alternate presentation.

**Figure 6.**
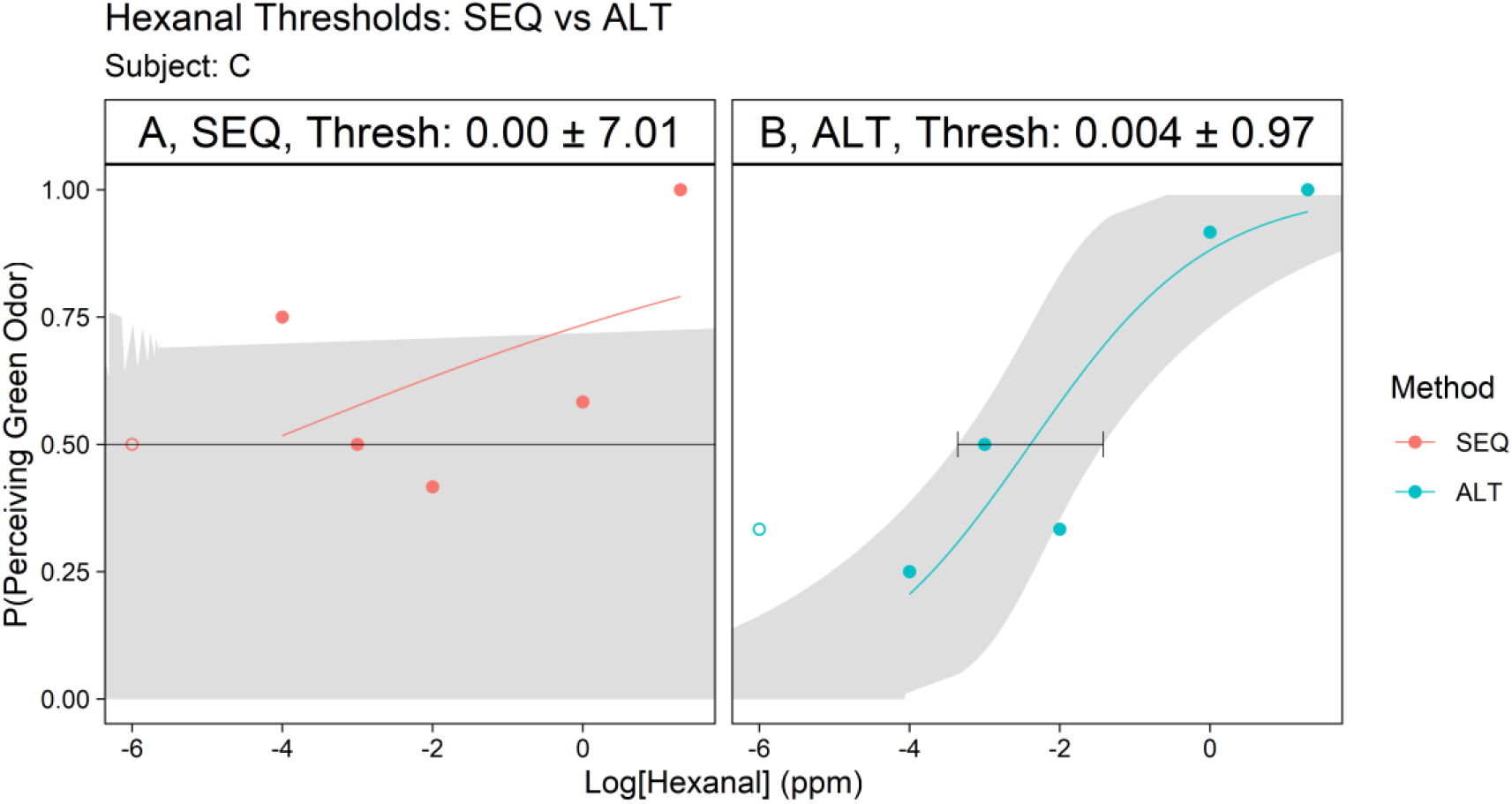
HEX threshold measurement with a blank sample (no odorant), subject C, replication 2. The samples were freshly made. The x-axis is the log10 of the sample concentrations (in ppm), and the y-axis is the probability the subject perceived a sample and identified it as “green”. For HEX, 15, 10, 5, 1, 0.1, and 0.01 ppm and for TMP, 20, 10, 5, 1, 0.5, and 0.1 ppm were chosen. The hollow point is when the blank sample (0 ppm) was instead of the lowest concentration.

Across all 7 subjects and across both stimulants (Hex and TMP), the ALT method predicted threshold values with smaller confidence intervals than the SEQ presentation. Whisker plots combining all the data (Figure 7) shows the greater precision of the alternate procedure when compared to the sequential procedure.

**Figure 7.**
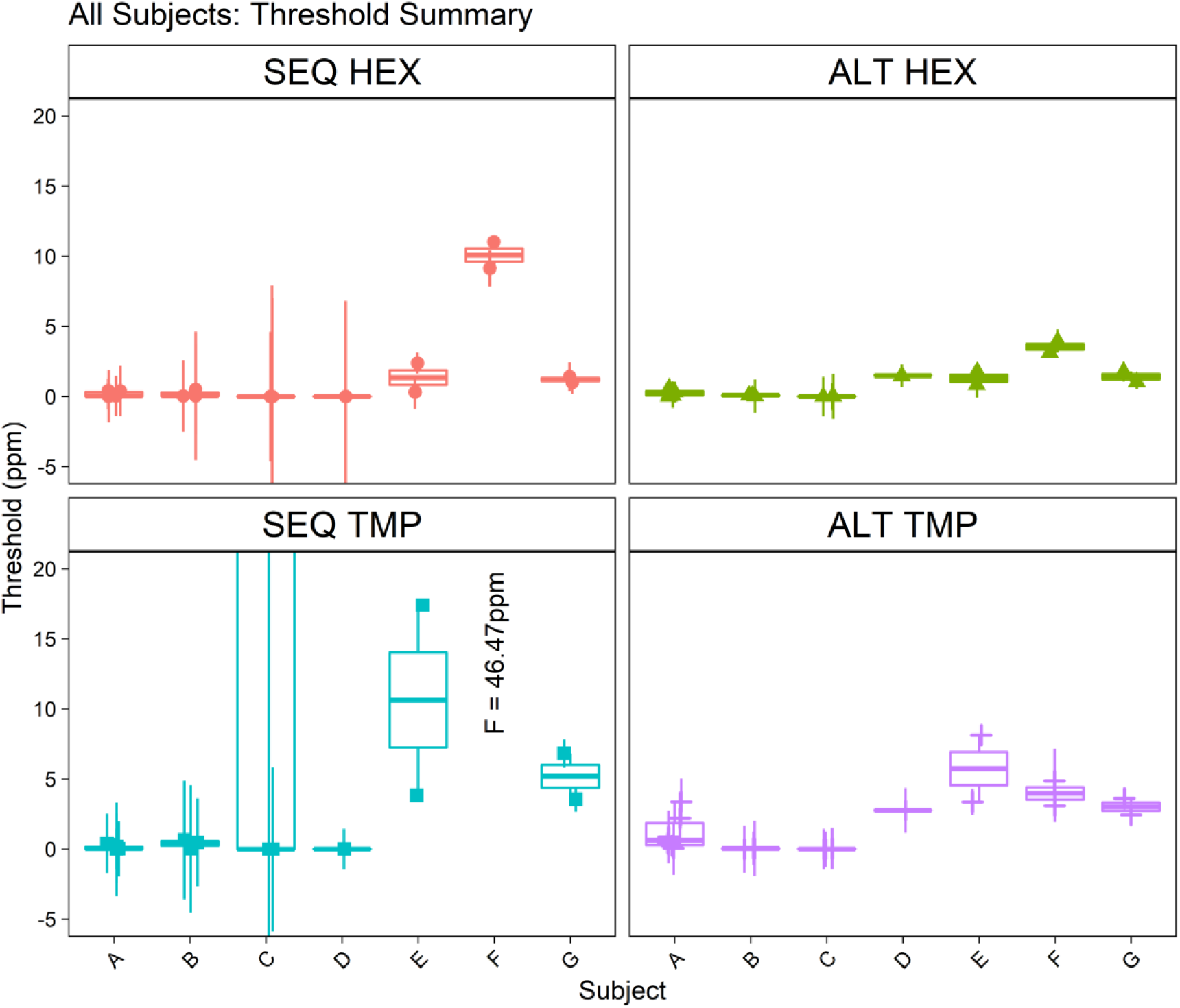
SEQ and ALT Boxplots for both HEX (15, 10, 5, 1, 0.1, and 0.01 ppm) and TMP (20, 10, 5, 1, 0.5, and 0.1 ppm) for all the subjects in all experiments. Each point represents the threshold for the subject on one test day using fresh stimulants, the boxplots are a compilation measurement of multiple test days.

### The phantosmia of blanks (No Odorant)

Phantosmia, sometimes called “odor hallucinations” is the recognition of an odor when no odorant is present(Greenberg 1992). Phantom odors have been reported by patients with schizophrenia, depression, and eating disorders(Kopala *et al*. 1994).In the thresholds where subjects were provided with a 10% PEG blank, all subjects (A, B, and C) tended to respond with “green” or “nutty” when the blank sample was presented.

As is shown in Figure 8, the probability a subject detected an odor (a phantom or ‘hallucination’) when smelling a blank ranged from p = 0.00 to p = 0.92. However, when smelling the lowest concentration—indicated by the level of the grey bar, an average measurement of detecting the lowest concentration for those subjects—, the probability they detected an odor was less than 0.15. Taken together, these experiments predict that the most robust psychophysical functions require interlaced sample presentations like the ALT method, and the absence of blanks containing no odorants.

**Figure 8.**
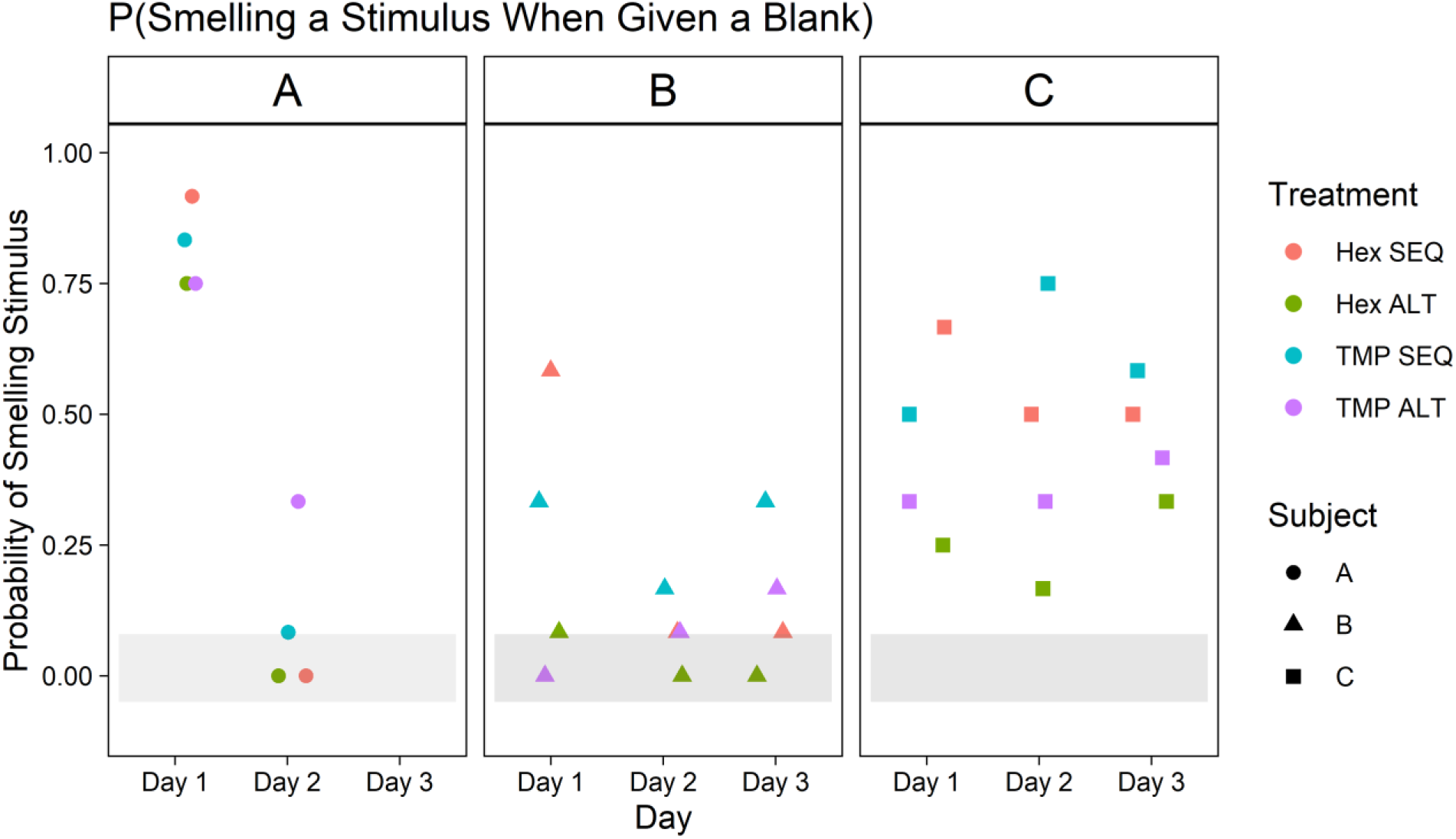
Response Probability for subjects A, B, and C when a blank sample (no odorant, 0 ppm) was presented instead of the lowest concentration. All the responses to the lowest concentration (0.001 and 0.0001 ppm) fell within the grey bar when no blank was presented.

## Discussion

It may seem counterintuitive that the ALT presentation of stimuli produced thresholds with lower variance and higher replicability than the SEQ presentation unless there was an interaction between adjacent puffs of stimuli during a session. The stimulus triads that made up a session were randomly sampled 12 times each while occupying all three positions in the sample block. The difference in concentration between adjacent puffs during the SEQ presentation was **Δ**x and 2**Δ**x while the adjacent pairs in the ALT presentation differed by 2**Δ**x and 4**Δ**x.

One possibility is that the concentration difference between adjacent pairs of puffs in a SEQ triad were sometimes below the JND (Just Noticeable Difference) hypothesized by Weber, while this happened less often during the ALT experiments. A calculation of the Weber fractions, **Δ**r/r, where **Δ**r is the difference in concentration between consecutive puffs and r is the concentration of the first puff ranged between 0.75 and 1.00. The Weber Ratios, the **Δ**r/r when the **Δ**r = the JND for 83 odorous substances averaged 0.25 to 0.33^32^, indicating that all the pairwise presentations in the ALT method were well above the JND(Egert *et al*. 2013). The Weber Ratio is a constant for each stimulus domain, dependent on the JND for that domain, below which two stimuli are indistinguishable. By contrast, the Weber fraction is the relationship between physical values of two stimuli. However, the Weber fractions of the binary pairs used in the SEQ method ranged from 0.25 and 0.80 indicating that some of the pairs may not be distinguishable. If the Weber fractions are indeed involved in creating the difference between the performance of the SEQ and ALT presentations, then the odor-perceptual process is based on sensations that are contiguous, distinguishable, and recognizable.

The inability to recognize a blank in the context of varying concentrations of a conditioned olfactory “space” would explain its random assignments; the blank does not fit into the olfactory space defined by the conditioning. By comparison, the ALT method provides complete coverage of the conditioned olfactory “space” with any given triad, covering both extremes of detection. This allows the production of a space wherein a subsequent test of a different—but interlacing—triad could fit quantitively in the same space, creating the precise, reproducible psychophysical curves obtained with the ALT method. Mathematically, we can define the “image” of a psychophysical function as its domain (the set of all probabilities produced by the set of all stimulants) and the image of the threshold concentration is the 0.5 probability of recognizing “grass” or “nuts” but only in the context of the conditioning and the presentation procedure.

To understand the theoretical implications of the results reported here requires further experiments (ex; varying the time between triads or between trials) to reveal more about this perceptual space and the structure of the “image” it contains. Recent work conducted by Schoonover et al on representational drift in the piriform cortex of mice(Schoonover *et al*. 2021) may lend support to the concept of olfactory identity being linked to relationships between stimuli, rather than absolute values of concentration. If identity is a property of noticeable difference between stimuli, a product of differences within the perceptual space, an odor identity may remain robust as long as a noticeable difference between stimuli is maintained.

The results of the experiments not only reveal procedural requirements for robust SO experiments, but additionally indicate that binary comparisons of brief odorant puffs may be perceived as differences between puffs, and not as the value of the puff itself. These findings may have further implications for the nature of how the brain processes and interprets olfactory stimuli, in addition to better informing how to measure and interrogate these processes.

## Tables

**Table 1.** Comparison between ALT and SEQ presentation methods for HEX (hexanal).

| Median Threshold (ppm) | Range | Method | Subject | Chemical |
| --- | --- | --- | --- | --- |
| 0.0395 | 0.415 | SEQ | A | Hexanal |
| 0.047 | 0.486 | SEQ | B | Hexanal |
| 0 | 0.001 | SEQ | C | Hexanal |
| 0.003 | 0 | SEQ | D | Hexanal |
| 1.352 | 2.072 | SEQ | E | Hexanal |
| 10.0895 | 1.893 | SEQ | F | Hexanal |
| 1.2125 | 0.421 | SEQ | G | Hexanal |
| 0.2355 | 0.592 | ALT | A | Hexanal |
| 0.098 | 0.073 | ALT | B | Hexanal |
| 0.004 | 0.003 | ALT | C | Hexanal |
| 1.491 | 0 | ALT | D | Hexanal |
| 1.3075 | 0.949 | ALT | E | Hexanal |
| 3.55 | 0.826 | ALT | F | Hexanal |
| 1.42 | 0.736 | ALT | G | Hexanal |

**Table 2.**
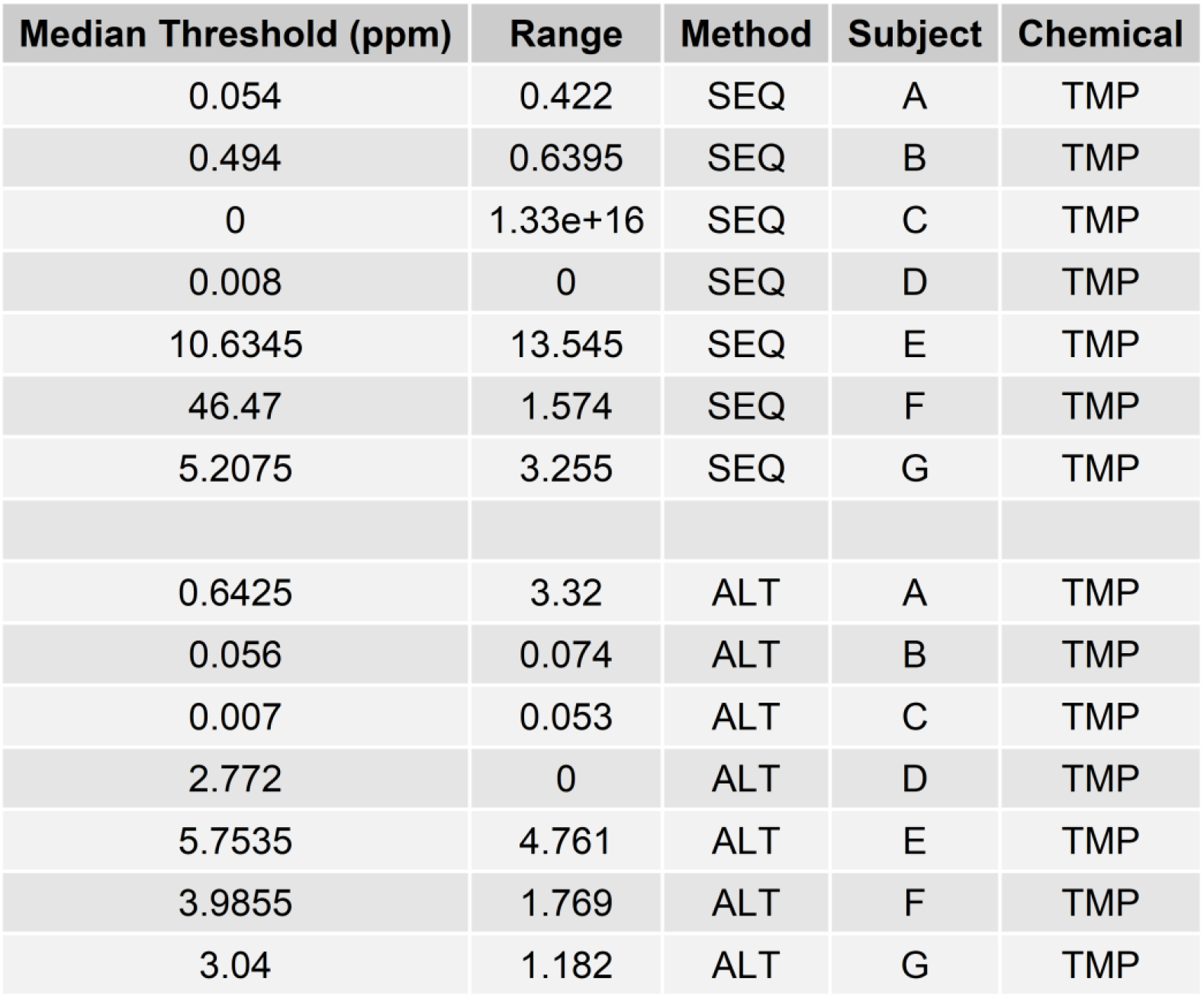
Comparison between ALT and SEQ presentation methods for TMP (2,2,3-trimethylpyrazine).

| Median Threshold (ppm) | Range | Method | Subject | Chemical |
| --- | --- | --- | --- | --- |
| 0.054 | 0.422 | SEQ | A | TMP |
| 0.494 | 0.6395 | SEQ | B | TMP |
| 0 | 1.33e+16 | SEQ | C | TMP |
| 0.008 | 0 | SEQ | D | TMP |
| 10.6345 | 13.545 | SEQ | E | TMP |
| 46.47 | 1.574 | SEQ | F | TMP |
| 5.2075 | 3.255 | SEQ | G | TMP |
| 0.6425 | 3.32 | ALT | A | TMP |
| 0.056 | 0.074 | ALT | B | TMP |
| 0.007 | 0.053 | ALT | C | TMP |
| 2.772 | 0 | ALT | D | TMP |
| 5.7535 | 4.761 | ALT | E | TMP |
| 3.9855 | 1.769 | ALT | F | TMP |
| 3.04 | 1.182 | ALT | G | TMP |

## Notes

https://github.com/terryacree/Ni22

### Competing Interest Statement

The authors have declared no competing interest.

## References

Uncategorized References

Barwich A-S. 2014. A Sense So Rare: Measuring Olfactory Experiences and Making a Case for a Process Perspective on Sensory Perception. Biological Theory 9: 258–268.

Cain W. 1979. To know with the nose: keys to odor identification. Science 203: 467–470.

Cain WS, Algom D. 1997. Perceptual and mental mixtures in odor and in taste: Are there similarities and differences between experiments or between modalities? Reply. J Exp Psychol Human 23: 1588–1593.

Cornsweet TN. 1962. The Staircase-Method in Psychophysics. The American Journal of Psychology 75: 485–491.

Delahunty CM, Eyres G, Dufour J-P. 2006. Gas chromatography-olfactometry. Journal of Separation Science 29: 2107–2125.

Ding K, Wang2 X, Tang Q, Albietz C, Rinberg D, Acree T. 2019. Sniff Olfactometry:Temporal effects on odorant mixture perception in humans. Chemarxiv.

Dravnieks A. 1975. Instrumental Aspects of Olfactometry. In: Moulton DG, Turk A, Johnson Jr. JW, (eds.), Methods in Olfactory Research. London: Academic Press. p. 1–51.

Egert M, Hohne HM, Weber T, Simmering R, Banowski B, Breves R. 2013. Identification of compounds inhibiting the C-S lyase activity of a cell extract from a Staphylococcus sp. isolated from human skin. Lett Appl Microbiol 57: 534–539.

Foster D, Scofield EH, Dallenbach KM. 1950. An Olfactorium. The American Journal of Psychology 63: 431–440.

Goyert HF, Frank ME, Gent JF, Hettinger TP. 2007. Corrigendum to “Characteristic component odors emerge from mixtures after selective adaptation” [Brain Res. Bull. 72 (2007) 1–9]. Brain Research Bulletin 73: 330.

Greenberg M. 1992. Olfactory Hallucinations. In: Serby MJ, Chobor KL, (eds.), Science of Olfaction. New York, NY: Springer.

Herz RS, Larsson M, Trujillo R, Casola MC, Ahmed FK, Lipe S, Brashear ME. 2022. A three-factor benefits framework for understanding consumer preference for scented household products: psychological interactions and implications for future development. Cogn Res Princ Implic 7: 28.

Kopala LC, Goos KP, Honer WG. 1994. Olfactory hallucinations and olfactory identification ability in patients with schizophrenia and other psychiatric disorders. Schizophrenia Research 12: 205–211.

Laing DG, Eddy A, Francis GW, Stephens L. 1994. Evidence for the temporal processing of odor mixtures in humans. Brain Research 651: 317–328.

Lawless HT, Heymann H. 2010. Sensory Evaluation of Food. New York: Springer New York.

Leland JV, Schieberle P, Buettner A, Acree TE, Editors. 2001. Gas chromatography-olfactometry: The state of the art. (Proceedings of a Symposium held at the 219th ACS National Meeting in New Orleans, Louisiana during August of 1999.) [In: ACS Symp. Ser., 2001; 782]. ACS.

MASS SFaDfVaRs. 2020. Modern Applied Statistics with S. CRAN.

Peirce J. 2022. PsychoPy.

Peirce J, Gray JR, Simpson S, MacAskill M, Hochenberger R, Sogo H, Kastman E, Lindelov JK. 2019. PsychoPy2: Experiments in behavior made easy. Behav Res Methods 51: 195–203.

Peirce JW. 2007. Psychophysics software in Python. Journal of Neuroscience Methods 162: 8–13.

Pellegrino R, Sinding C, de Wijk RA, Hummel T. 2017. Habituation and adaptation to odors in humans. Physiol Behav 177: 13–19.

Rankin CH. 2009. Introduction to special issue of neurobiology of learning and memory on habituation. Neurobiol Learn Mem 92: 125–126.

Rochelle MM. 2017. The Psychophysical Perception of the Key Odorants in Potato Chips. Food Science. Ithaca, NY: Cornell University.

Rochelle MM, Prévost GA, Acree TE. 2017a. Computing Odorant Images. J Agric Food Chem.

Rochelle MM, Prévost GJ, Acree TE. 2017b. Computing Odor Images. Journal of Agricultural and Food Chemistry 66: 2219–2225.

Romagny S, Coureaud G, Thomas-Danguin T. 2018. Key odorants or key associations? Insights into elemental and configural odour processing. Flavour and Fragrance Journal 33: 97–105.

Schmidt R, Cain WS. 2010. Making scents: dynamic olfactometry for threshold measurement. Chem Senses 35: 109–120.

Schoonover CE, Ohashi SN, Axel R, Fink AJP. 2021. Representational drift in primary olfactory cortex. Nature 594: 541–546.

Sinding C, Thomas-Danguin T, Chambault A, Béno N, Dosne T, Chabanet C, Schaal B, Coureaud G. 2013. Rabbit Neonates and Human Adults Perceive a Blending 6-Component Odor Mixture in a Comparable Manner. PLOS ONE 8: e53534.

Sinding C, Thomas-Danguin T, Crepeaux G, Schaal B, Coureaud G. 2011. Experience influences elemental and configural perception of certain binary odour mixtures in newborn rabbits. The Journal of Experimental Biology 214: 4171–4178.

Team RC. 2020. R: A language and environment for statistical computin g. Vienna, Austria.

Thomas-Danguin T, Sinding C, Romagny S, El Mountassir F, Atanasova B, Le Berre E, Le Bon AM, Coureaud G. 2014. The perception of odor objects in everyday life: a review on the processing of odor mixtures. Front Psychol 5: 504.

Wilson DA, Linster C. 2008. Neurobiology of a simple memory. J Neurophysiol 100: 2–7.

Wilson DA, Stevenson RJ. 2006. Learning to Smell: Olfactory Perception from Neurobiology to Behavior. Baltimore: The Johns Hopkins University Press.

Wise PM, Olsson MJ, Cain WS. 2000. Quantification of Odor Quality. Chem Senses 25: 429–443.

Wyckoff SG, Acree TE. 2017. Multi-Modal Olfactometer - US Patent 9,642,570. In: Office USP, editor Us Patent Application Publication. USA: Stephen Wyckoff.

Zhao H, Ivic L, Otaki JM, Hashimoto M, Mikoshiba K, Firestein S. 1998. Functional Expression of a Mammalian Odorant Receptor. Science 279: 241–242.

