## Supplemental Materials and Documents for "‘Sniff Olfactometer (SO) Protocols"

Supplemental Documentation & Figures

### Code and Analysis

For all R and PsychoPy code used, as well as the Cornell IRB notice, please visit our GitHub repository here: https://github.com/terryacree/Ni22

### Research Participant Information and Consent Form

**Description of the research**

You are invited to participate in a research study about olfactory perception. The participation is voluntary. The purpose of this study is to investigate olfactory perception, and establish new protocols for doing so.

**Confidentiality**

This study is anonymous. We will not be collecting or retaining any information about your identity. The record of this study will be kept strictly confidential. Research records will be kept in a locked file, and all electronic information will be coded and secured using a password-protected file. Your identity will not be disclosed in the material that is published.

**What will my participation involve?**

In the research, you will first be trained to be familiar with aromas of hexanal (green) and 2,3,5-trimethylpyrazine (nut); all chemicals used in the study are food-grade. After the training, you will be tested on the identification of the same aromatic substances. If 87% correct score is achieved, you will move to the 2nd part of the study, or you have to retake the training session. The repetition will not exceed 5 times.

Before the 2nd part, you are allowed to take 5-10 minutes of rest. In the 2nd part, you will be asked to smell and identify puffs of odorants using the sniff olfactometer.

The experiment is a forced-choice experiment, so you will have to make a choice even if you are not sure about your answer. The total running time per session is about 15-30 minutes, and you are allowed to take up to 10-20 minutes of rest between any 2 sessions.

**Responsibilities**

If you decide to participate in this study, you will be asked to follow these items—

1. You are responsible NOT to apply any fragrance products (e.g., perfume).
2. You are responsible NOT to smoke, drink, or eat at least 1 hour before the study.
3. You are responsible for informing the researcher about your nose condition (e.g., stuffed nose).
4. You are responsible for wearing headphones/earphones throughout the entire experiment to eliminate experimental error (headphones and earphones can be prepared by yourself).
5. You are encouraged to ask questions and comment anytime during the experiment, but NOT all the questions will be answered.
6. You MAY be asked about your opinion about the entire experimental procedure.

**Compensation**

Upon proper completion of the study, you will receive a $10 reward.

**Who to contact**

You may present your questions to the experiment administrator Terry Acree at any time. The researcher conducting this study is Jiayue Ni. Please feel free to contact Jiayue Ni at. Your participation in this study is completely voluntary, and should you feel it necessary at any time to withdraw, alert the administrator. If you have any questions or concerns regarding your rights as a subject in this study, you may contact the Cornell Institutional Review Board (IRB) at 607-255-5138 or access their website at http://www.irb.cornell.edu. You may also report your concerns or complaints anonymously through Ethicpoints (www.hotline.cornell.edu) or by calling at 1-866-293-3077. Ethicpoints is an independent organization that serves as a liaison between the University and the person bringing any complaints so that anonymity can be ensured.

*By signing the consent form, I understand the purpose of this study and what I will be asked to do. I know the confidentiality will be ensured and I will receive proper compensation. I also am aware of my responsibilities and rights.*

Name: ___________________________ Date: __________________________

(Prepared originally and modified by the author with permission from Terry Acree)

### Supplemental Figures

### Subject A

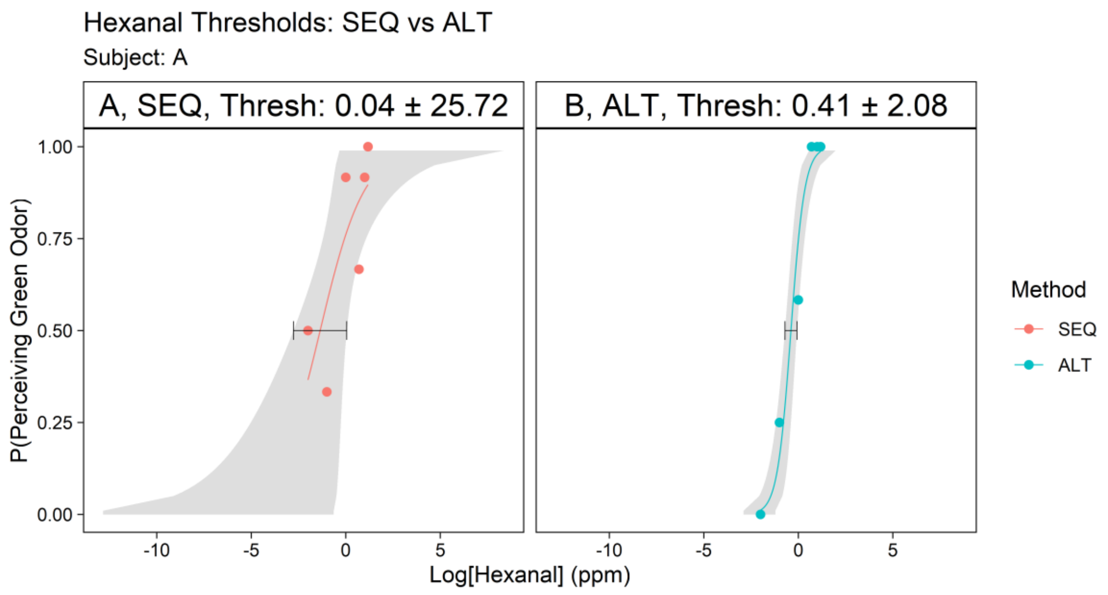


Figure A.1. Solution reproducibility test -- HEX threshold measurement, subject A, day 1. Solutions were freshly made. The x-axis is the log10 of the sample concentrations (in ppm), and the y-axis is the probability when the subject perceived a sample with the veridical "green".

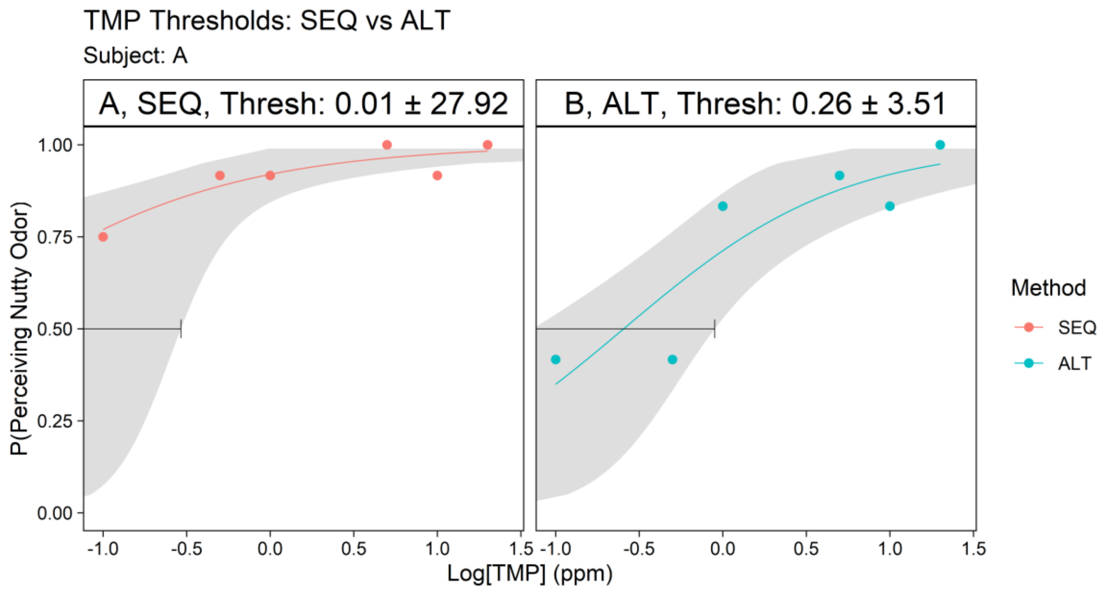


Figure A.2. Solution reproducibility test -- TMP threshold measurement, subject A, day 1. Solutions were freshly made. The x-axis is the log10 of the sample concentrations (in ppm), and the y-axis is the probability when the subject perceived a sample with the veridical “nutty”.

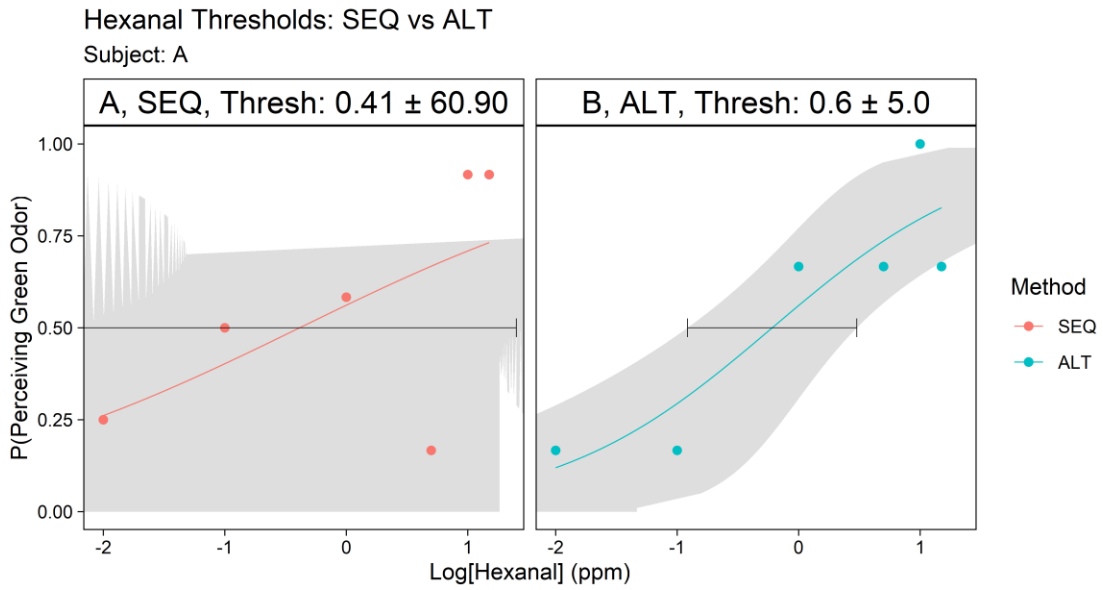


Figure A.3. Solution reproducibility test -- HEX threshold measurement, subject A, replication. Solutions were recycled from day 1. The x-axis is the log10 of the sample concentrations (in ppm), and the y-axis is the probability when the subject perceived a sample with the veridical “green”. The imprecision of the model confirmed the use of fresh samples for each run.

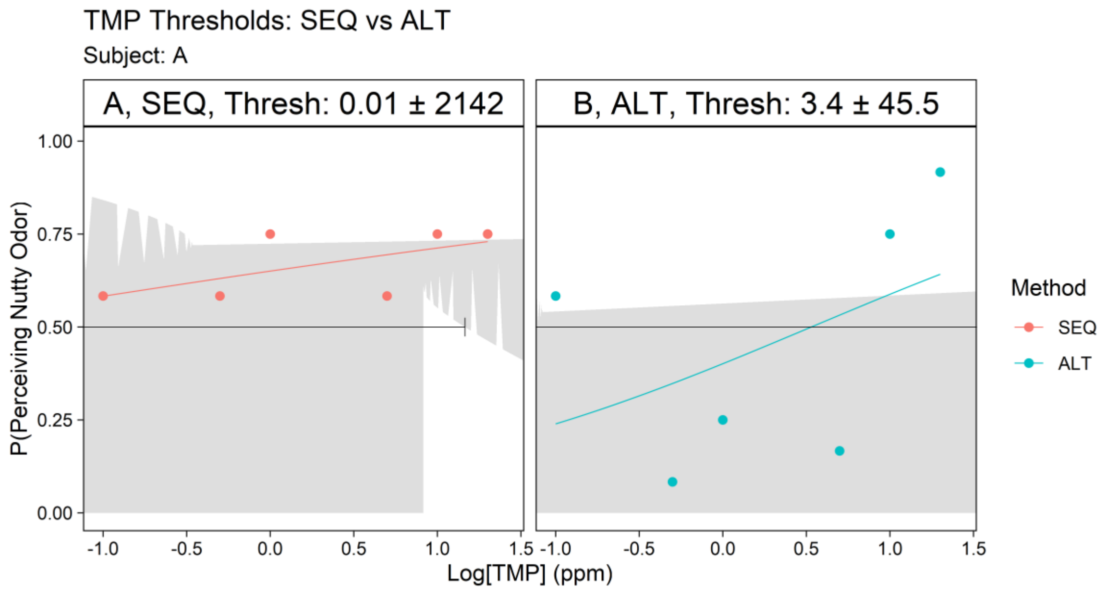


Figure A.4. Solution reproducibility test -- TMP threshold measurement, subject A, replication. Solutions were recycled from day 1. The x-axis is the log10 of the sample concentrations (in ppm), and the y-axis is the probability when the subject perceived a sample with the veridical "nutty". The imprecision of the model confirmed the use of fresh samples for each run.

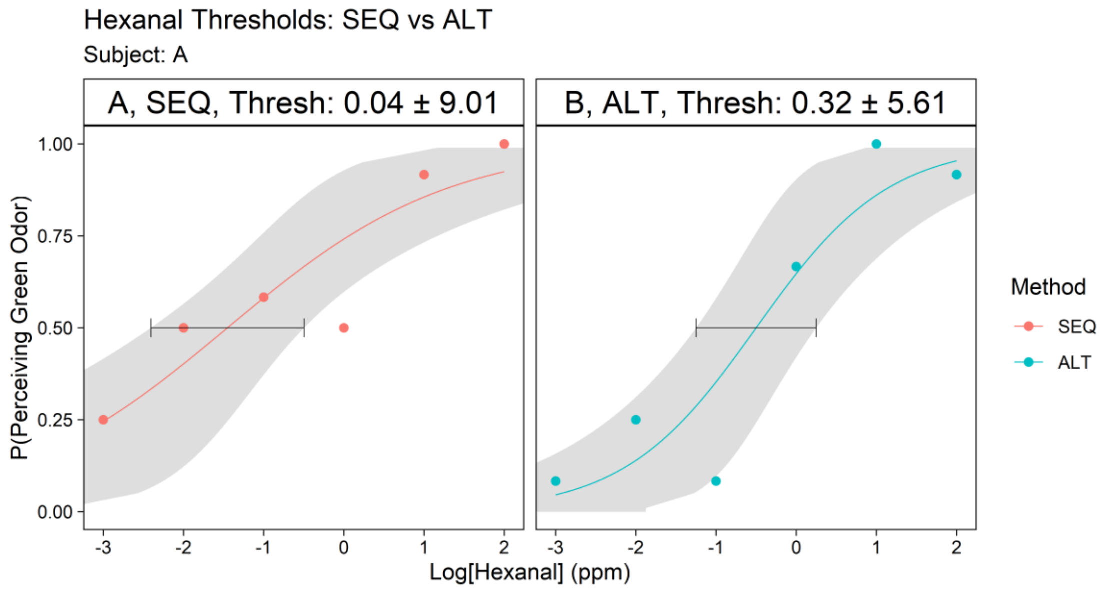


Figure A.5. Solution reproducibility test -- HEX threshold measurement with a high concentration (100 ppm), subject A, day 1. Solutions were freshly made. The x-axis is the log10 of the sample concentrations (in ppm), and the y-axis is the probability when the subject perceived a sample with the veridical "green".

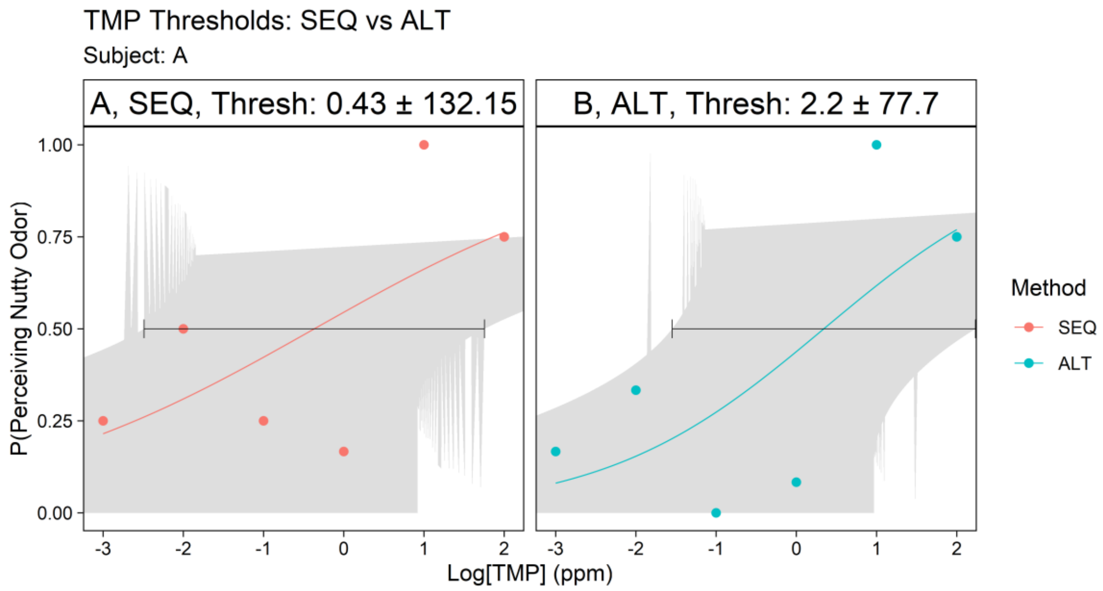


Figure A.6. Solution reproducibility test -- TMP threshold measurement with a high concentration (100 ppm), subject A, day 1. Solutions were freshly made. The x-axis is the log10 of the sample concentrations (in ppm), and the y-axis is the probability when the subject perceived a sample with the veridical “nutty”.

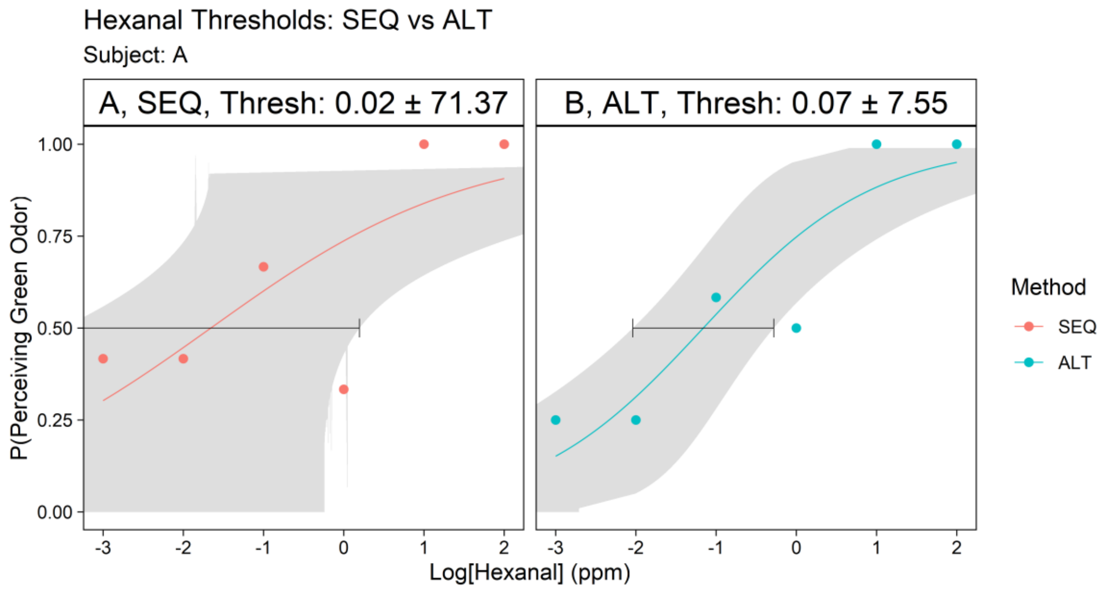


Figure A.7. Solution reproducibility test -- HEX threshold measurement with a high concentration (100 ppm), subject A, replication. Solutions were recycled from day 1. The x-axis is the log10 of the sample concentrations (in ppm), and the y-axis is the probability when the subject perceived a sample with the veridical “green”. The imprecision of the model confirmed the use of fresh samples for each run.

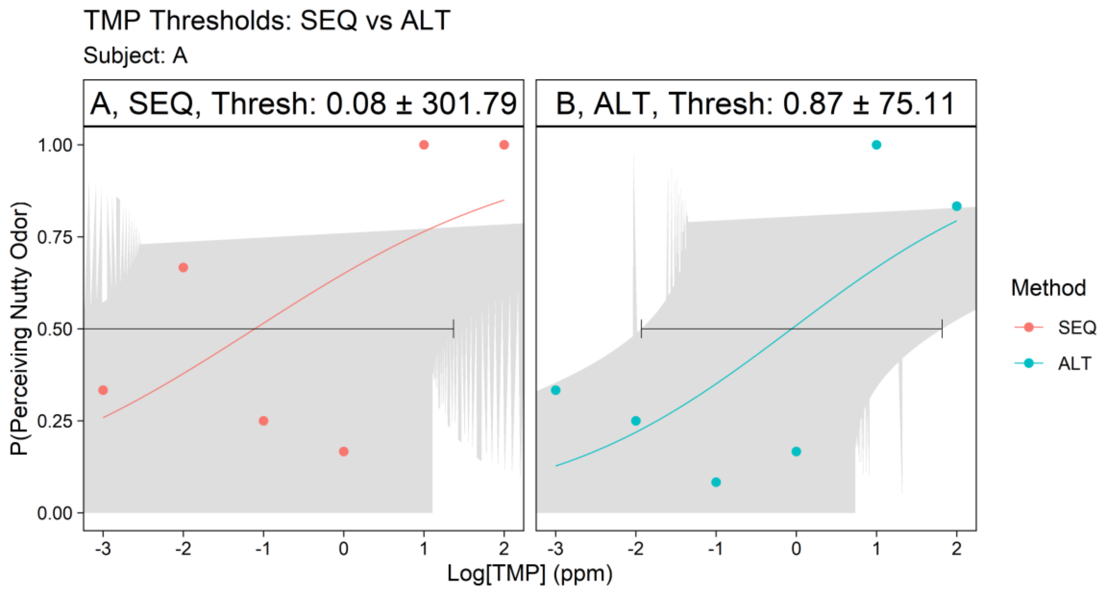


Figure A.8. Solution reproducibility test -- TMP threshold measurement with a high concentration (100 ppm), subject A, replication. Solutions were recycled from day 1. The x-axis is the log10 of the sample concentrations (in ppm), and the y-axis is the probability when the subject perceived a sample with the veridical "nutty". The imprecision of the model confirmed the use of fresh samples for each run.

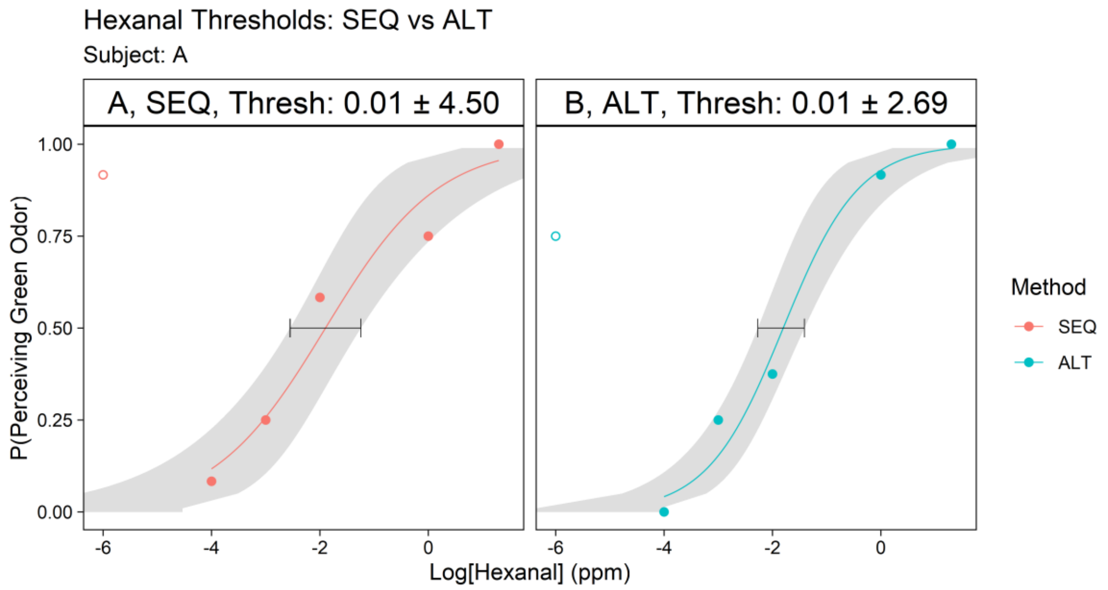


Figure A.9. Blank test -- HEX threshold measurement with a blank sample (no odorant), subject A, day 1. Solutions were freshly made. The hollow point is when the blank sample (0 ppm) was exposed to the subject. The x-axis is the log10 of the sample concentrations (in ppm), and the y-axis is the probability when the subject perceived a sample with the veridical "green".

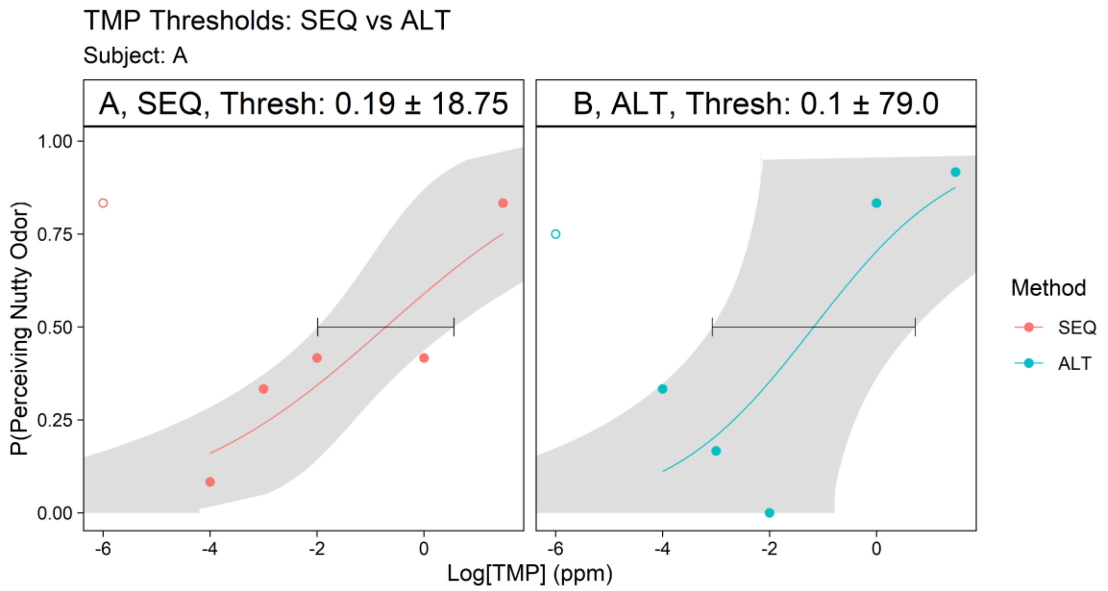


Figure A.10. Blank test -- TMP threshold measurement with a blank sample (no odorant), subject A, day 1. Solutions were freshly made. The hollow point is when the blank sample (0 ppm) was exposed to the subject. The x-axis is the log10 of the sample concentrations (in ppm), and the y-axis is the probability when the subject perceived a sample with the veridical "nutty".

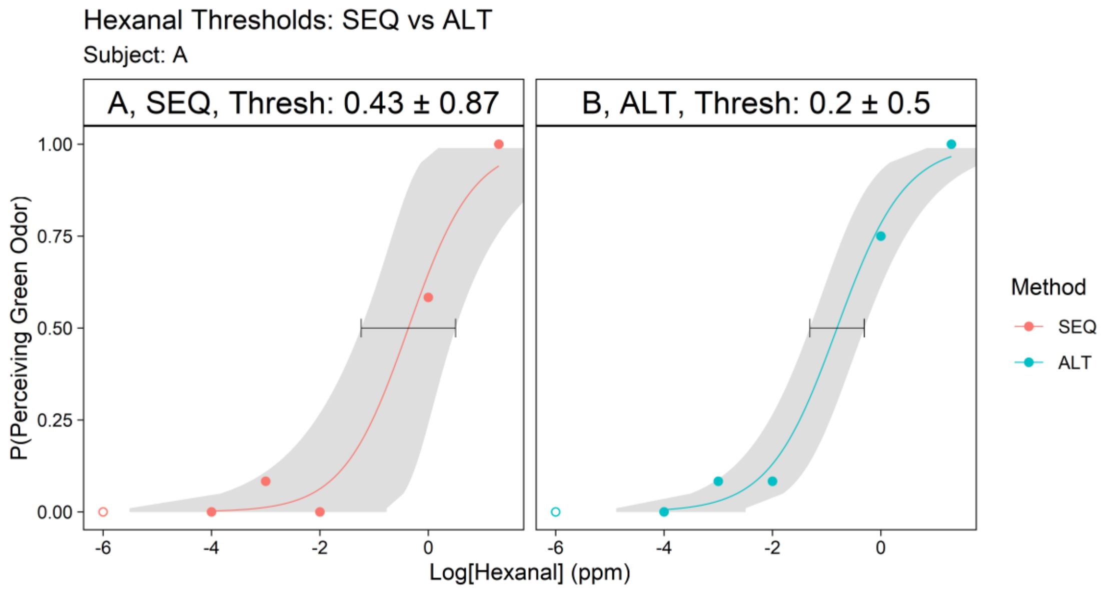


Figure A.11. Blank test -- HEX threshold measurement with a blank sample (no odorant), subject A, day 2. Solutions were freshly made. The hollow point is when the blank sample (0 ppm) was exposed to the subject. The x-axis is the log10 of the sample concentrations (in ppm), and the y-axis is the probability when the subject perceived a sample with the veridical "green".

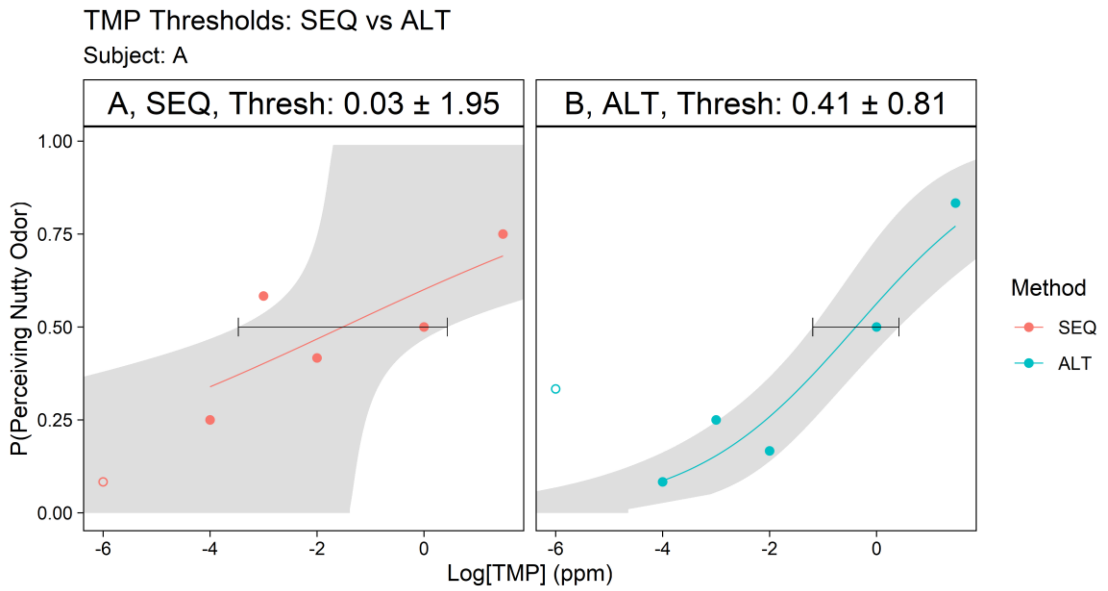


Figure A.12. Blank test -- TMP threshold measurement with a blank sample (no odorant), subject A, day 2. Solutions were freshly made. The hollow point is when the blank sample (0 ppm) was exposed to the subject. The x-axis is the log10 of the sample concentrations (in ppm), and the y-axis is the probability when the subject perceived a sample with the veridical “nutty”.

### Subject B

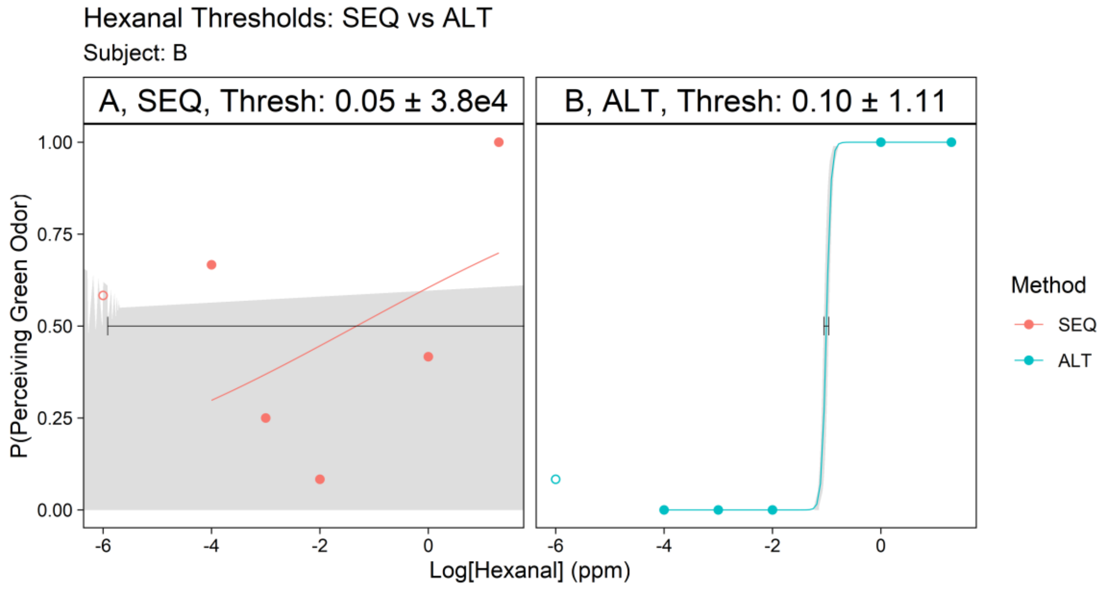


Figure B.1. HEX threshold measurement with a blank sample (no odorant), subject B, day 1. The hollow point is when the blank sample (0 ppm) was exposed to the subject. The x-axis is the log10 of the sample concentrations (in ppm), and the y-axis is the probability when the subject perceived a sample with the veridical “green”.

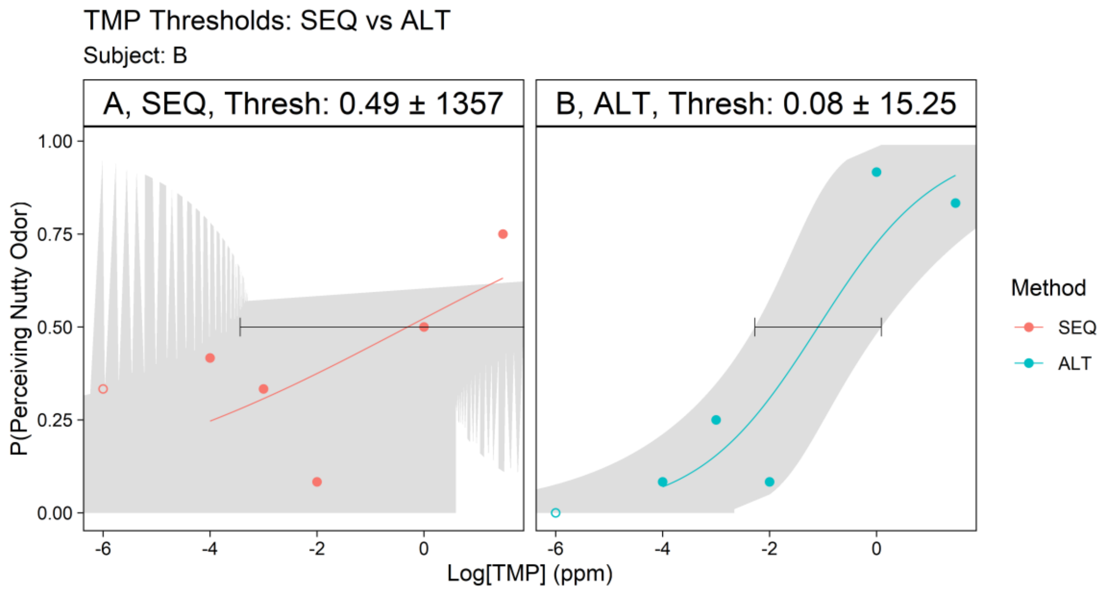


Figure B.2. TMP threshold measurement with a blank sample (no odorant), subject B, day 1. The hollow point is when the blank sample (0 ppm) was exposed to the subject. The x-axis is the log10 of the sample concentrations (in ppm), and the y-axis is the probability when the subject perceived a sample with the veridical "nutty".

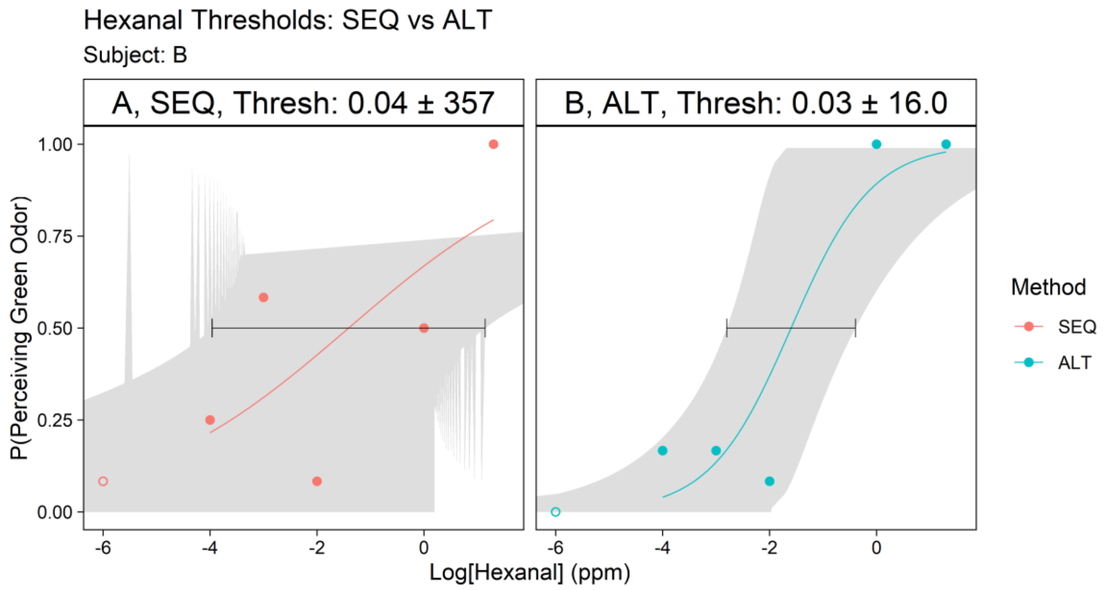


Figure B.3. HEX threshold measurement with a blank sample (no odorant), subject B, day 2. The hollow point is when the blank sample (0 ppm) was exposed to the subject. The x-axis is the log10 of the sample concentrations (in ppm), and the y-axis is the probability when the subject perceived a sample with the veridical "green".

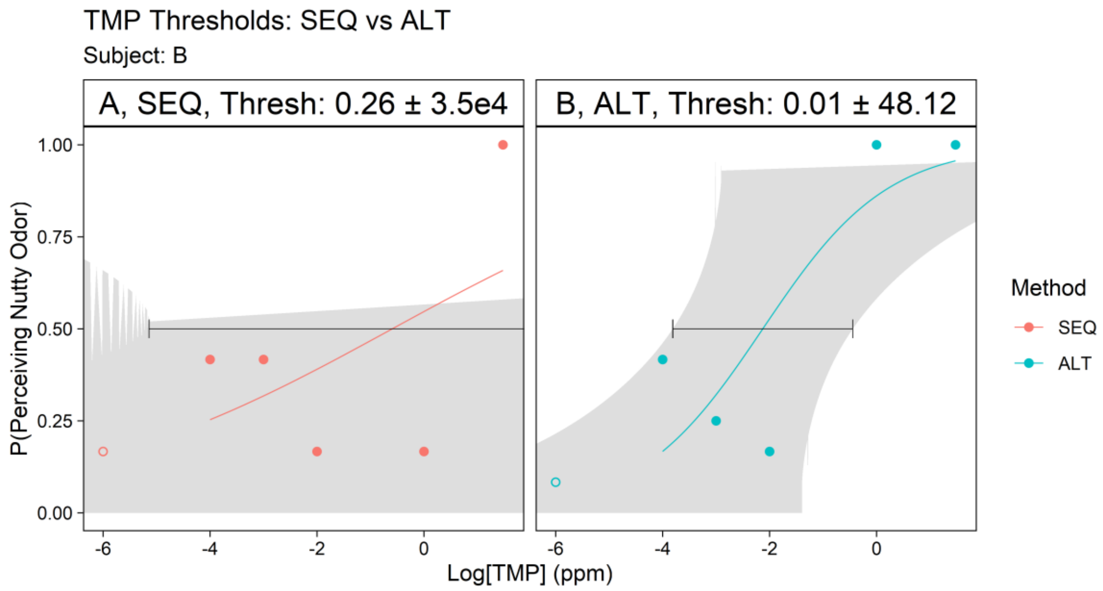


Figure B.4. TMP threshold measurement with a blank sample (no odorant), subject B, day 2. The hollow point is when the blank sample (0 ppm) was exposed to the subject. The x-axis is the log10 of the sample concentrations (in ppm), and the y-axis is the probability when the subject perceived a sample with the veridical “nutty”.

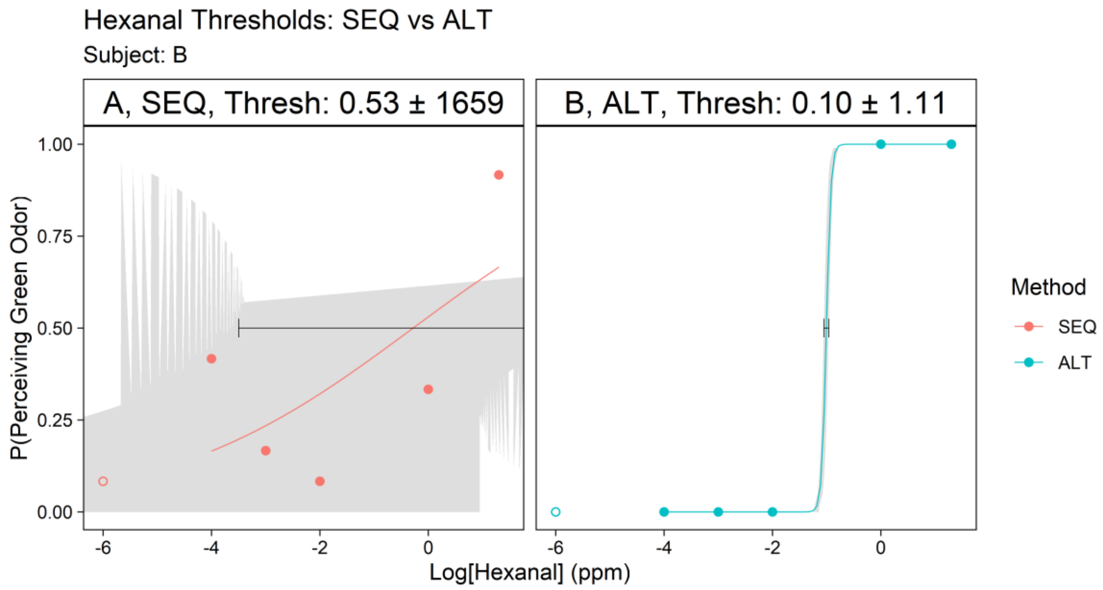


Figure B.5. HEX threshold measurement with a blank sample (no odorant), subject B, day 3. The hollow point is when the blank sample (0 ppm) was exposed to the subject. The x-axis is the log10 of the sample concentrations (in ppm), and the y-axis is the probability when the subject perceived a sample with the veridical “green”.

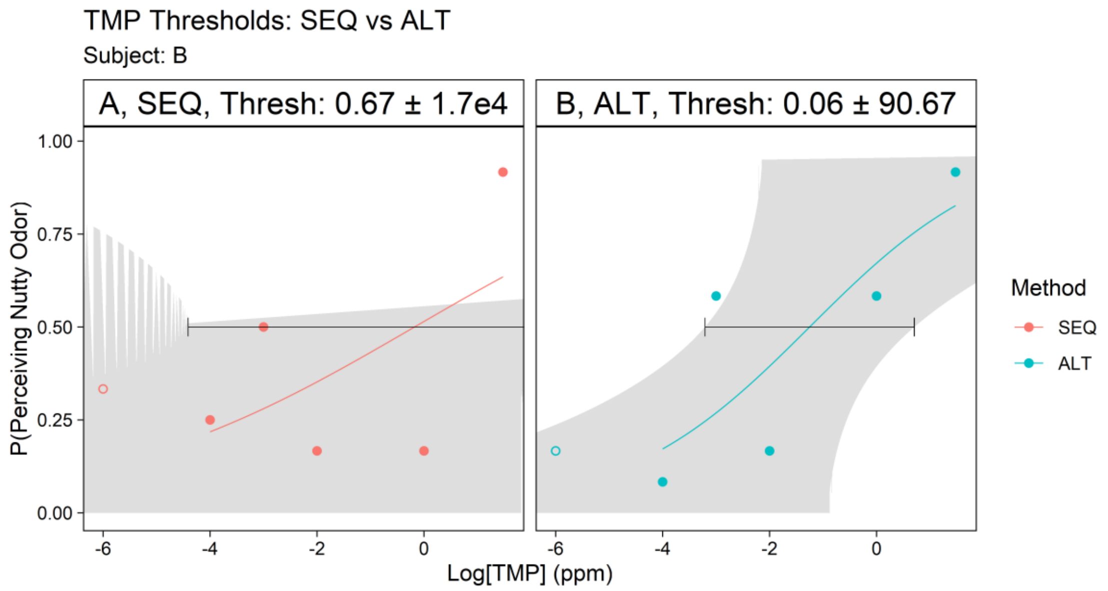


Figure B.6. TMP threshold measurement with a blank sample (no odorant), subject B, day 3. The hollow point is when the blank sample (0 ppm) was exposed to the subject. The x-axis is the log10 of the sample concentrations (in ppm), and the y-axis is the probability when the subject perceived a sample with the veridical “nutty”.

### Subject C

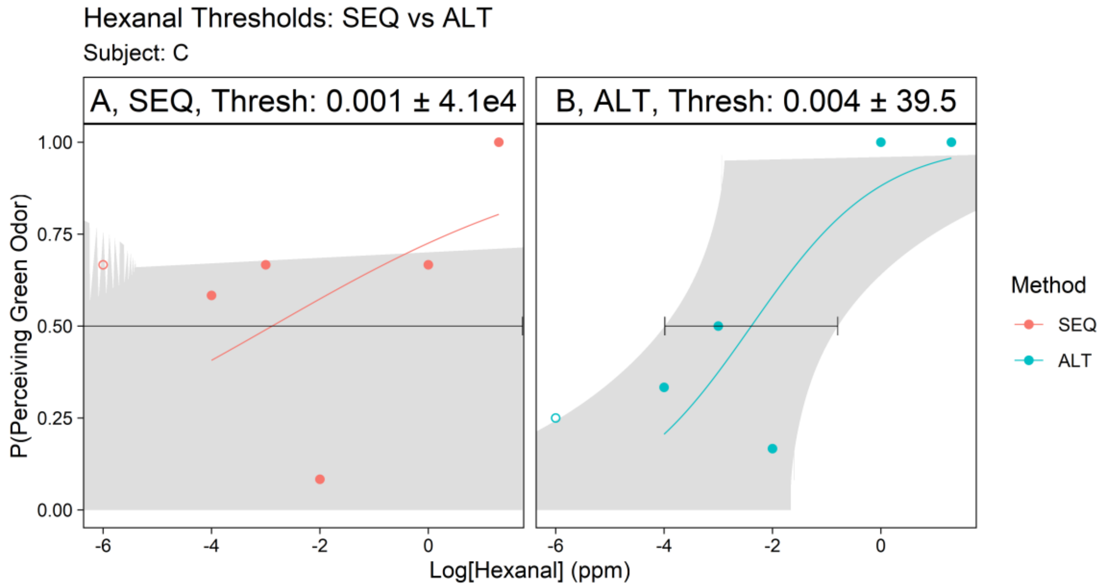


Figure C.1. HEX threshold measurement with a blank sample (no odorant), subject C, day 1. The hollow point is when the blank sample (0 ppm) was exposed to the subject. The x-axis is the log10 of the sample concentrations (in ppm), and the y-axis is the probability when the subject perceived a sample with the veridical “green”.

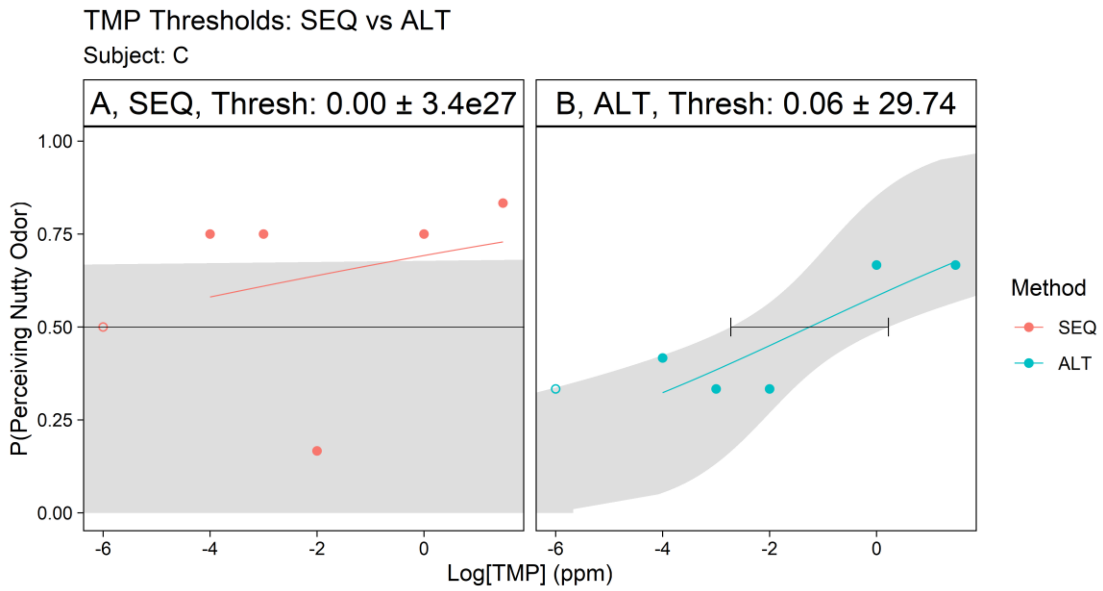


Figure C.2. TMP threshold measurement with a blank sample (no odorant), subject C, day 1. The hollow point is when the blank sample (0 ppm) was exposed to the subject. The x-axis is the log10 of the sample concentrations (in ppm), and the y-axis is the probability when the subject perceived a sample with the veridical “nutty”.

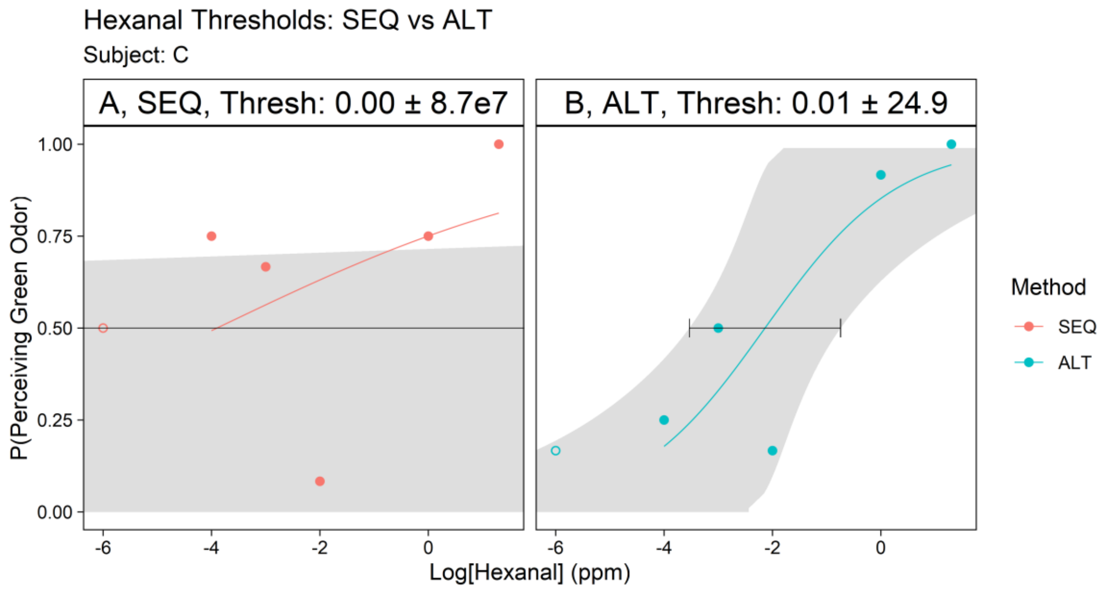


Figure C.3. HEX threshold measurement with a blank sample (no odorant), subject C, replication 1. The hollow point is when the blank sample (0 ppm) was exposed to the subject. The samples were freshly made. The x-axis is the log10 of the sample concentrations (in ppm), and the y-axis is the probability when the subject perceived a sample with the veridical “green”.

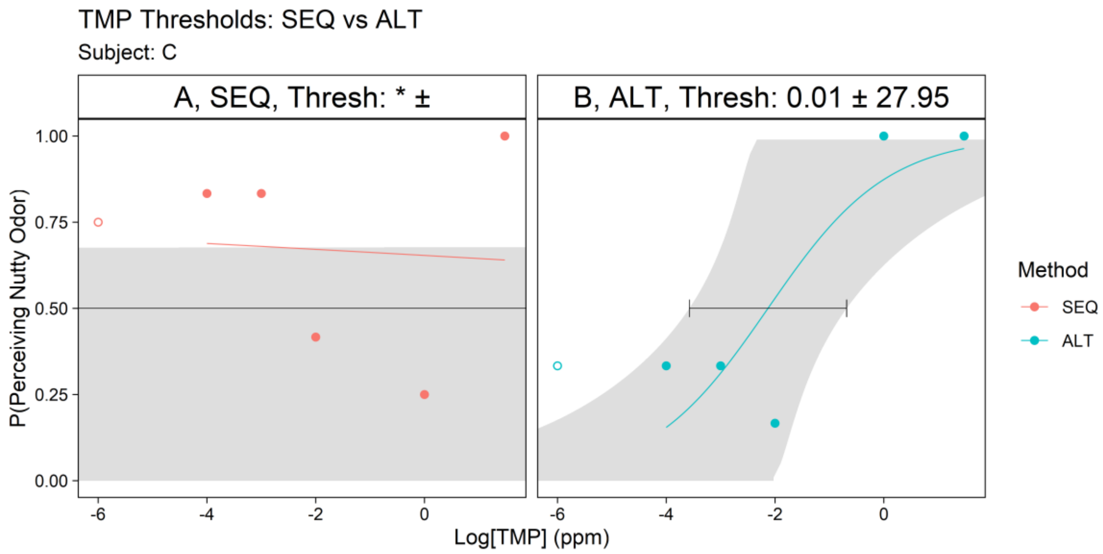


Figure C.4. TMP threshold measurement with a blank sample (no odorant), subject C, replication 1. The hollow point is when the blank sample (0 ppm) was exposed to the subject. The samples were freshly made. The x-axis is the log10 of the sample concentrations (in ppm), and the y-axis is the probability when the subject perceived a sample with the veridical “nutty”. * In this figure the calculated threshold was 1.3 × 1016 ppm, which is an unrealistic value. The possible explanation for that was that the subject was not able to systematically determine the concentrations in approximately their order of concentration with the SEQ order.

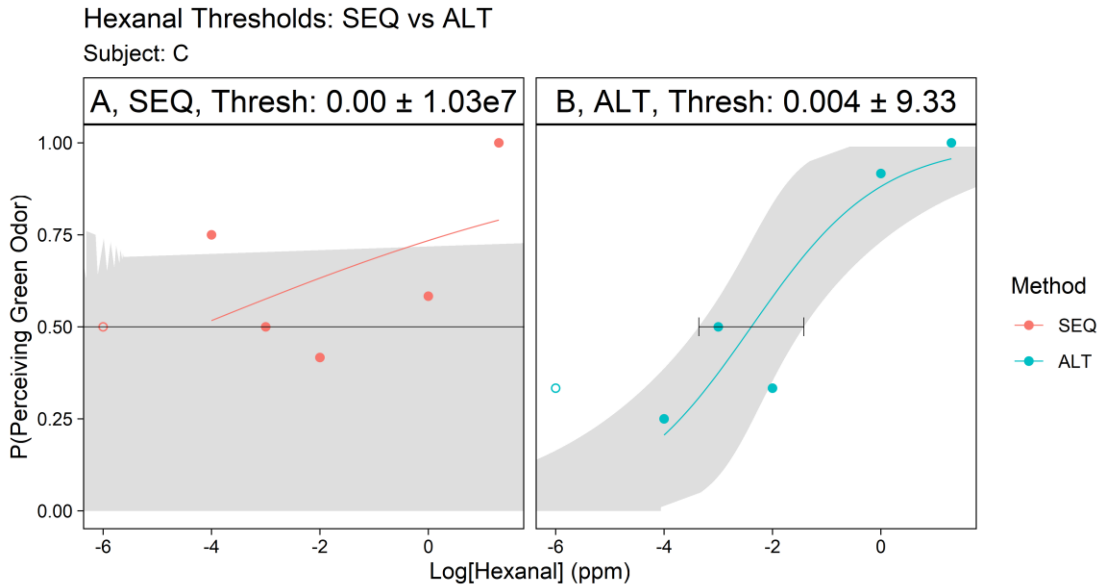


Figure C.5. HEX threshold measurement with a blank sample (no odorant), subject C, replication 2. The hollow point is when the blank sample (0 ppm) was exposed to the subject. The samples were freshly made. The x-axis is the log10 of the sample concentrations (in ppm), and the y-axis is the probability when the subject perceived a sample with the veridical “green”.

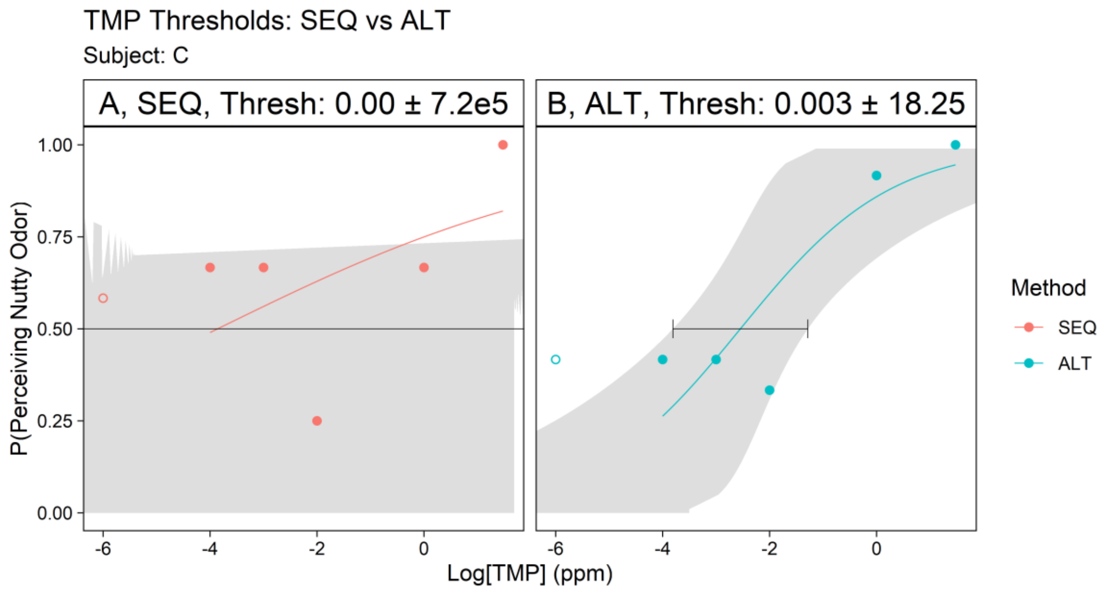


Figure C.6. TMP threshold measurement with a blank sample (no odorant), subject C, replication 2. The hollow point is when the blank sample (0 ppm) was exposed to the subject. The samples were freshly made. The x-axis is the log10 of the sample concentrations (in ppm), and the y-axis is the probability when the subject perceived a sample with the veridical "nutty".

### Subject D

This subject was only able to complete one day of testing.

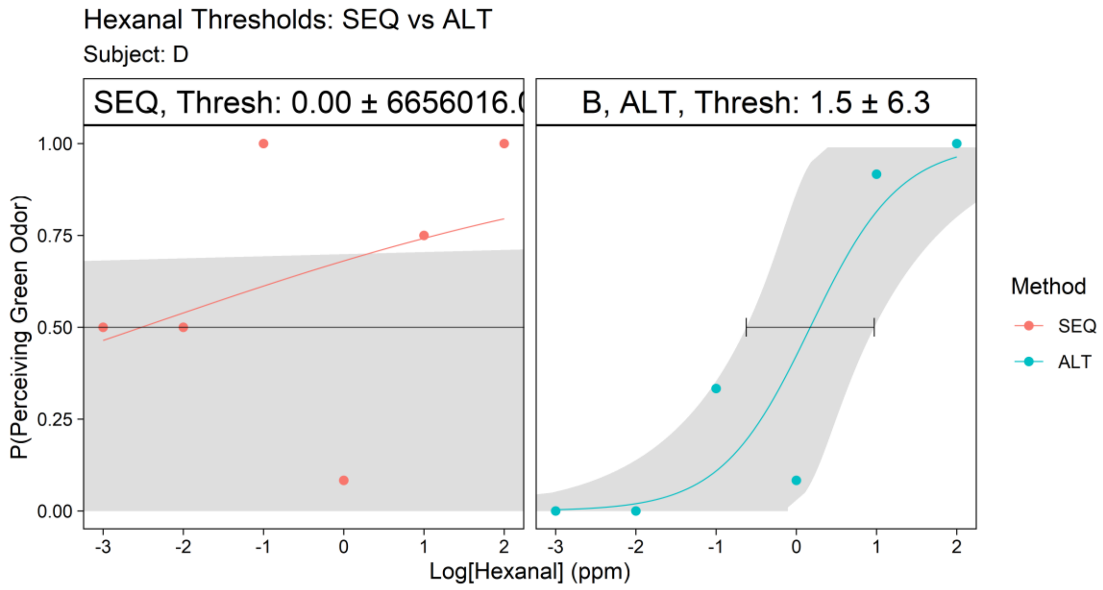


Figure D.1. HEX threshold measurement with a higher concentration (100 ppm), subject D, day 1. The samples were freshly made. The x-axis is the log10 of the sample concentrations (in ppm), and the y-axis is the probability when the subject perceived a sample with the veridical "green".

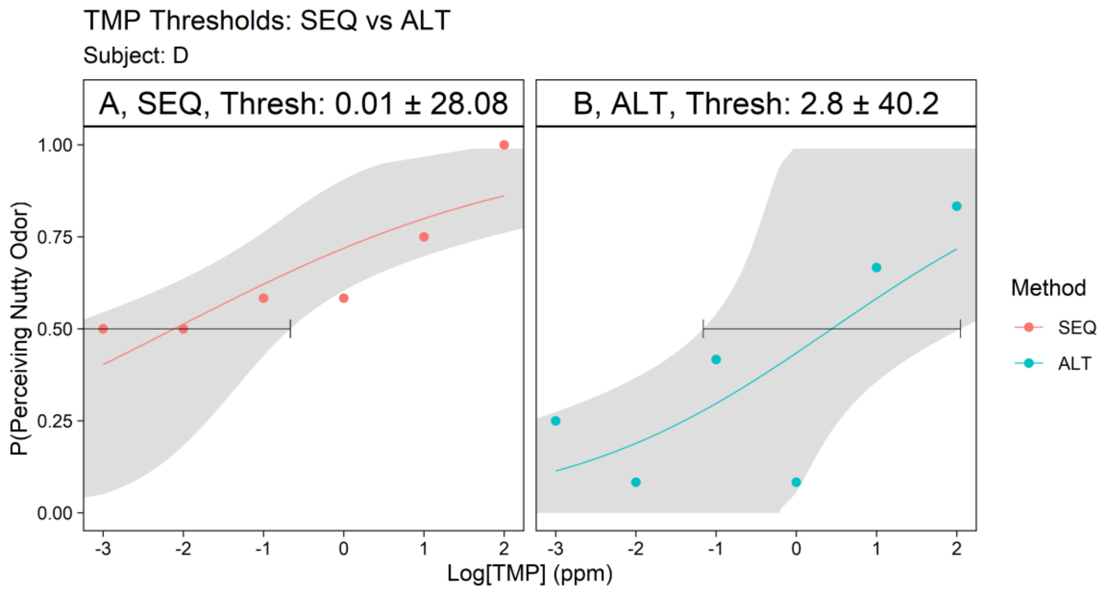


Figure D.2. TMP threshold measurement with a higher concentration (100 ppm), subject D, day 1. The x-axis is the log10 of the sample concentrations (in ppm), and the y-axis is the probability when the subject perceived a sample with the veridical "nutty".

### Subject E

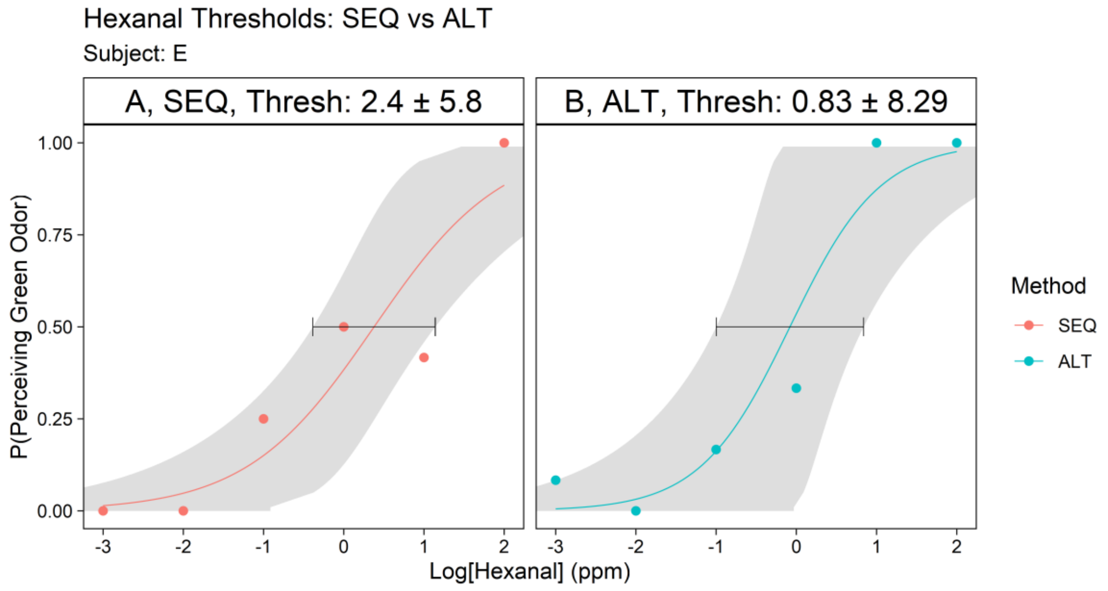


Figure E.1. HEX threshold measurement with a higher concentration (100 ppm), subject E, day 1. The samples were freshly made. The x-axis is the log10 of the sample concentrations (in ppm), and the y-axis is the probability when the subject perceived a sample with the veridical "green".

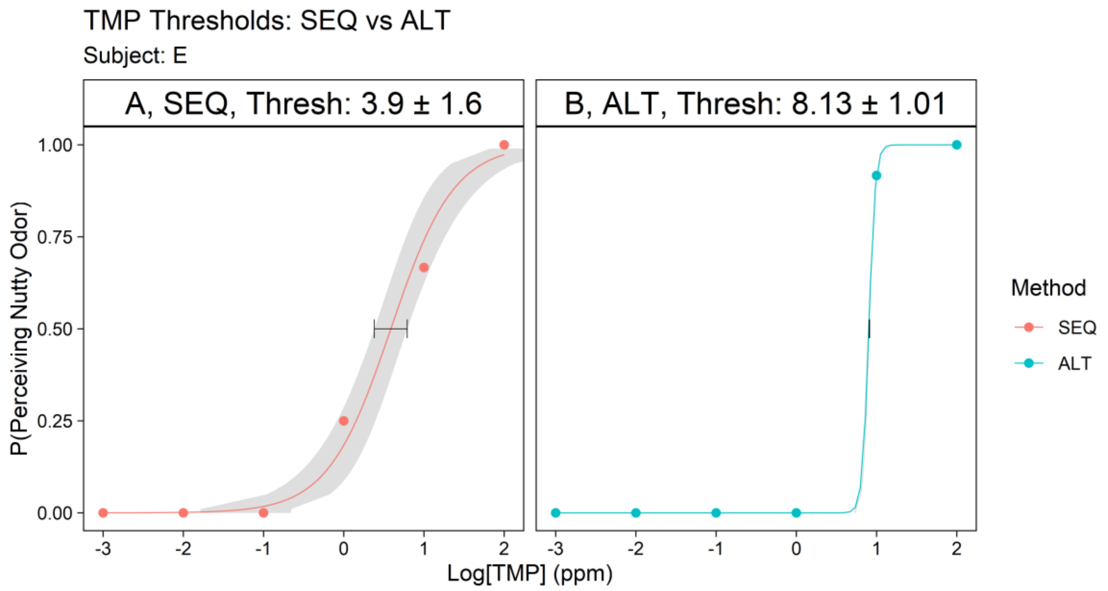


Figure E.2. TMP threshold measurement with a higher concentration (100 ppm), subject E, day 1. The samples were freshly made. The x-axis is the log10 of the sample concentrations (in ppm), and the y-axis is the probability when the subject perceived a sample with the veridical “nutty”.

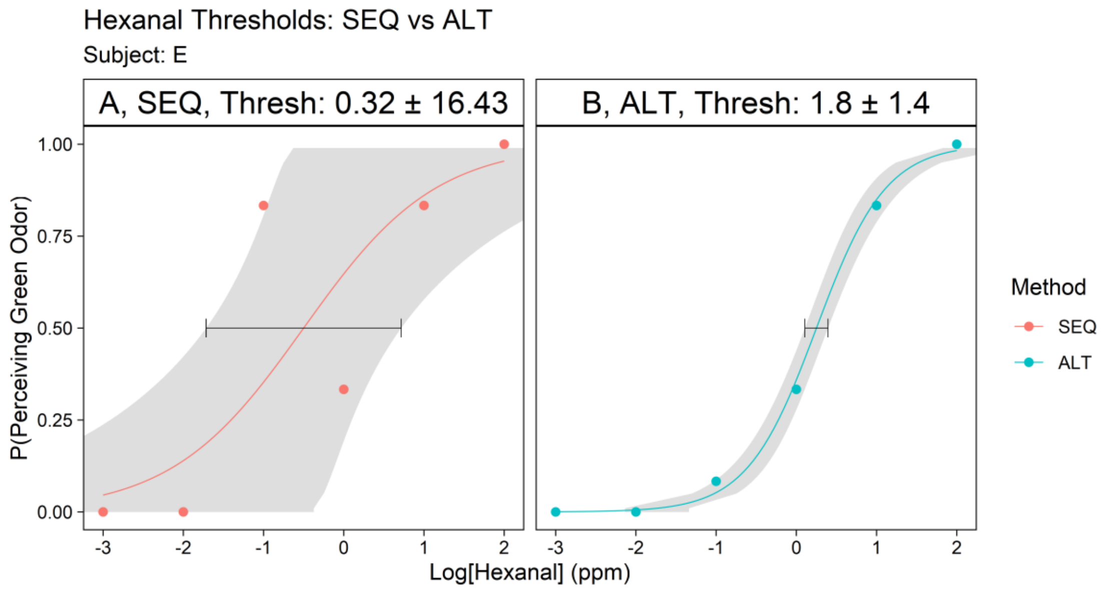


Figure E.3. HEX threshold measurement with a higher concentration (100 ppm), subject E, day 2. The samples were freshly made. The x-axis is the log10 of the sample concentrations (in ppm), and the y-axis is the probability when the subject perceived a sample with the veridical “green”.

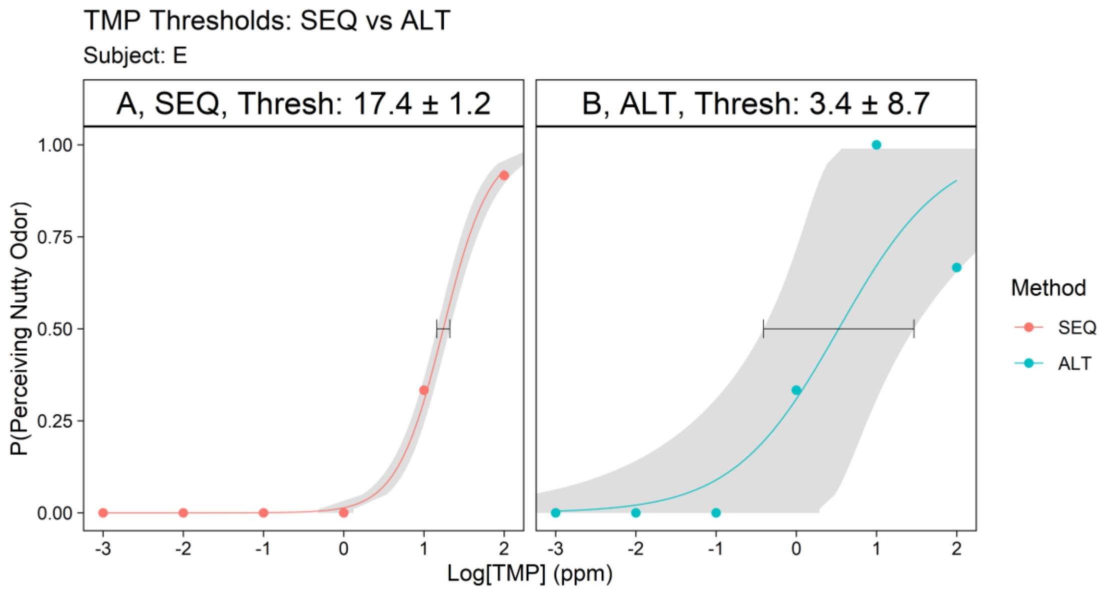


Figure E.4. TMP threshold measurement with a higher concentration (100 ppm), subject E, day 2. The samples were freshly made. The x-axis is the log10 of the sample concentrations (in ppm), and the y-axis is the probability when the subject perceived a sample with the veridical “nutty”.

### Subject F

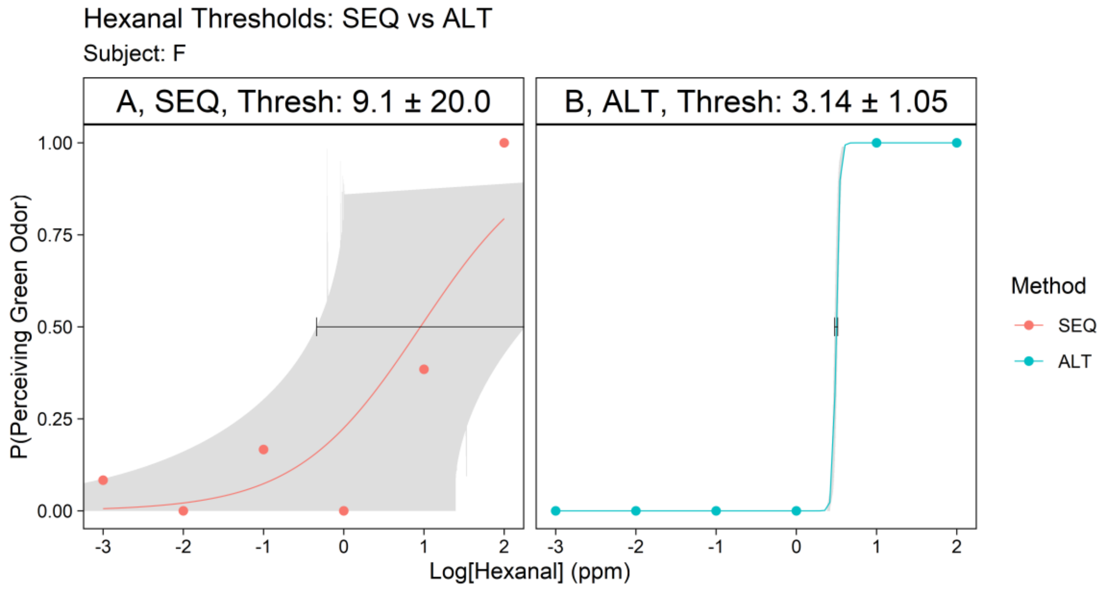


Figure F.1. HEX threshold measurement with a higher concentration (100 ppm), subject F, day 1. The samples were freshly made. The x-axis is the log10 of the sample concentrations (in ppm), and the y-axis is the probability when the subject perceived a sample with the veridical “green”.

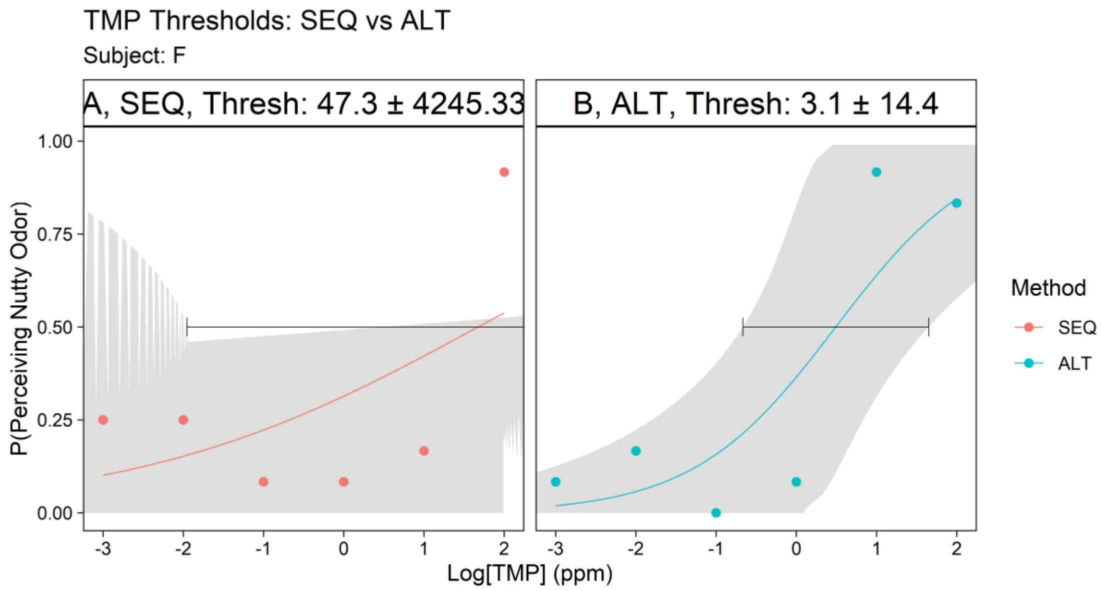


Figure F.2. TMP threshold measurement with a higher concentration (100 ppm), subject F, day 1. The samples were freshly made. The x-axis is the log10 of the sample concentrations (in ppm), and the y-axis is the probability when the subject perceived a sample with the veridical "nutty".

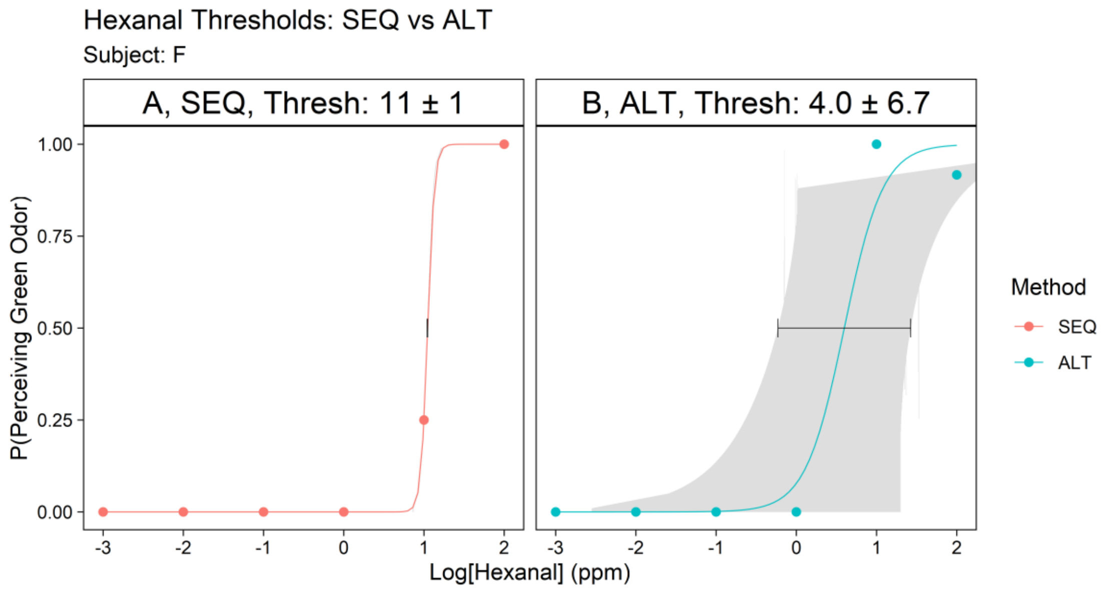


Figure F.3. HEX threshold measurement with a higher concentration (100 ppm), subject F, day 2. The samples were freshly made. The x-axis is the log10 of the sample concentrations (in ppm), and the y-axis is the probability when the subject perceived a sample with the veridical "green".

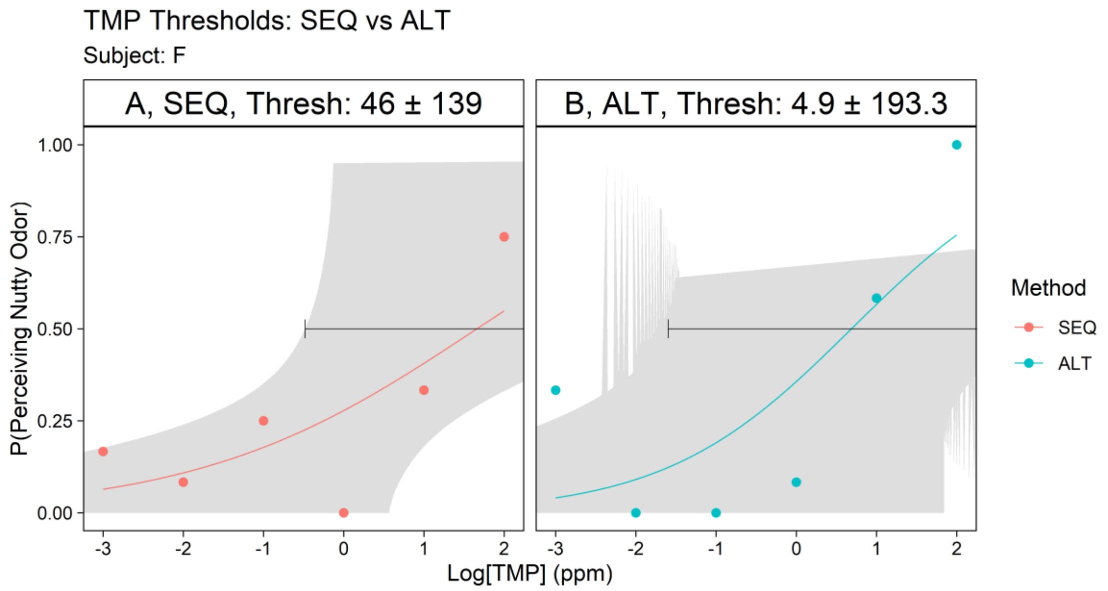


Figure F.4. TMP threshold measurement with a higher concentration (100 ppm), subject F, day 2. The samples were freshly made. The x-axis is the log10 of the sample concentrations (in ppm), and the y-axis is the probability when the subject perceived a sample with the veridical “nutty”.

### Subject G

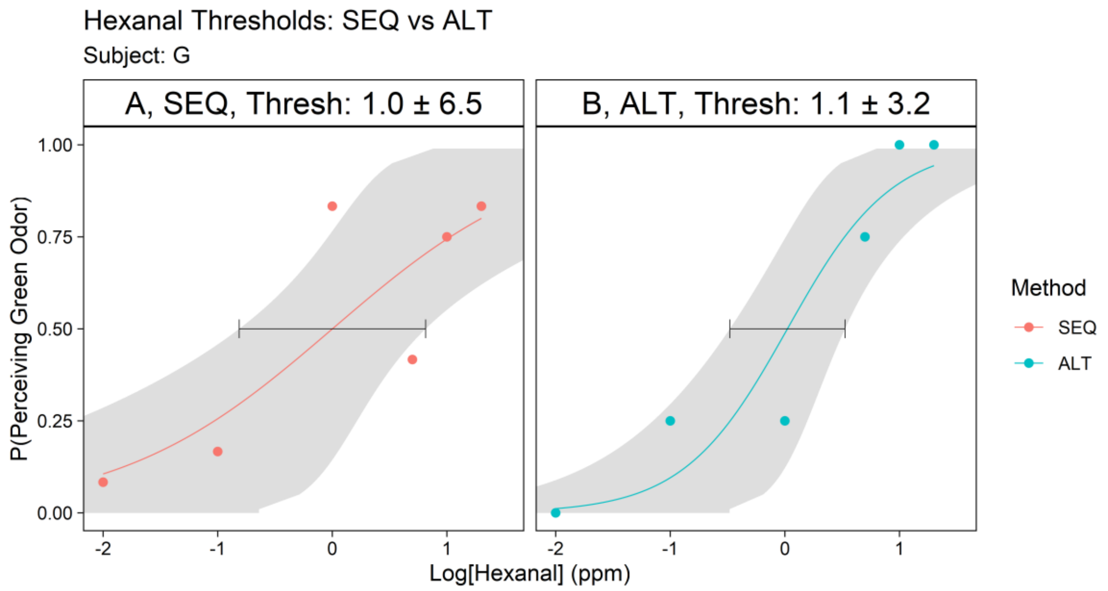


Figure G.1. HEX threshold measurement with a higher concentration (100 ppm), subject G, day 1. The samples were freshly made. The x-axis is the log10 of the sample concentrations (in ppm), and the y-axis is the probability when the subject perceived a sample with the veridical “green”.

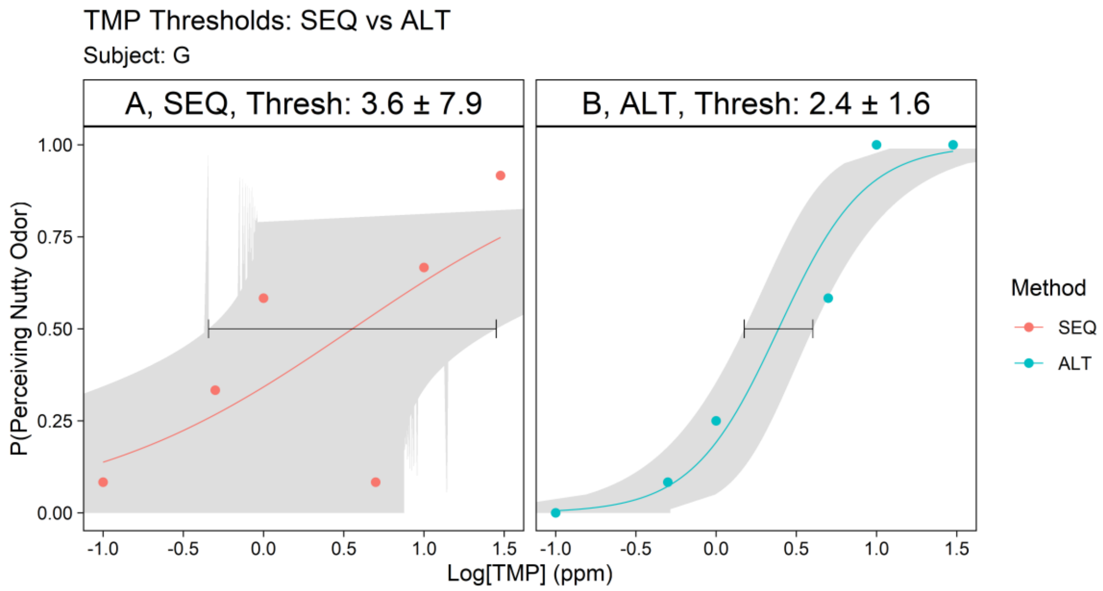


Figure G.2. TMP threshold measurement with a higher concentration (100 ppm), subject G, day 1. The samples were freshly made. The x-axis is the log10 of the sample concentrations (in ppm), and the y-axis is the probability when the subject perceived a sample with the veridical “nutty”.

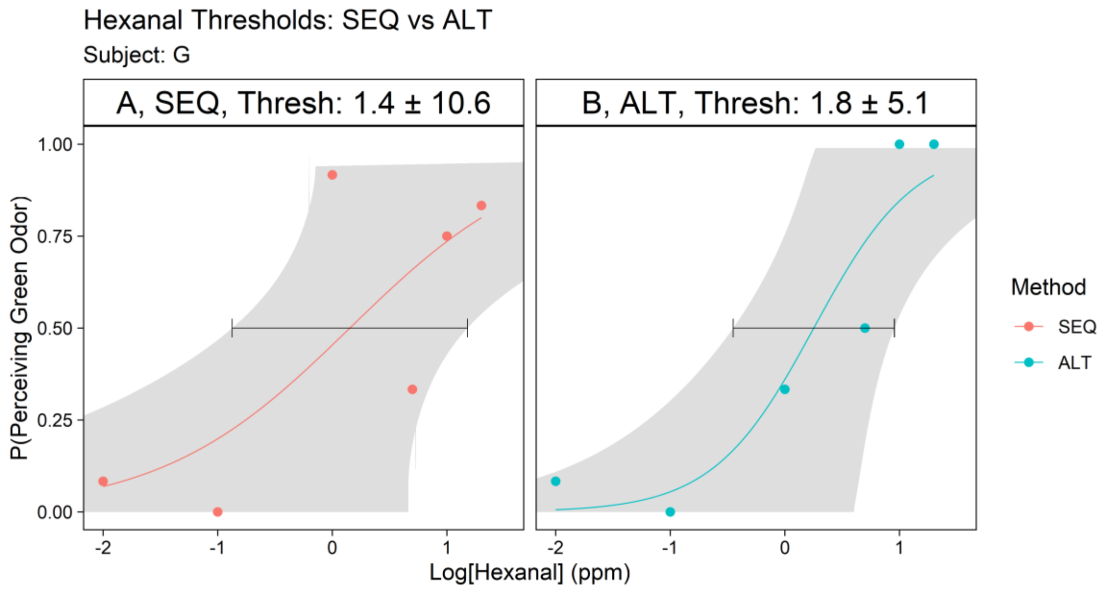


Figure G.3. HEX threshold measurement with a higher concentration (100 ppm), subject G, day 2. The samples were freshly made. The x-axis is the log10 of the sample concentrations (in ppm), and the y-axis is the probability when the subject perceived a sample with the veridical “green”.

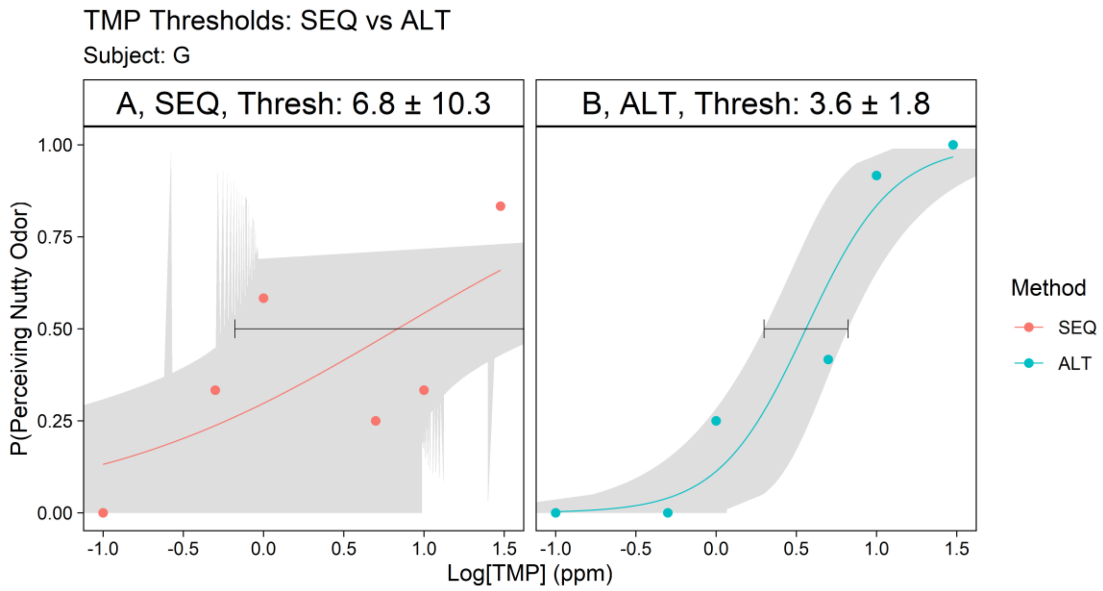


Figure G.4. TMP threshold measurement with a higher concentration (100 ppm), subject G, day 2. The samples were freshly made. The x-axis is the log10 of the sample concentrations (in ppm), and the y-axis is the probability when the subject perceived a sample with the veridical “nutty”.
